# Design of stimulus-responsive two-state hinge proteins

**DOI:** 10.1101/2023.01.27.525968

**Authors:** Florian Praetorius, Philip J. Y. Leung, Maxx H. Tessmer, Adam Broerman, Cullen Demakis, Acacia F. Dishman, Arvind Pillai, Abbas Idris, David Juergens, Justas Dauparas, Xinting Li, Paul M. Levine, Mila Lamb, Ryanne K. Ballard, Stacey R. Gerben, Hannah Nguyen, Alex Kang, Banumathi Sankaran, Asim K. Bera, Brian F. Volkman, Jeff Nivala, Stefan Stoll, David Baker

## Abstract

Proteins that switch between two structural states as a function of environmental stimuli are widespread in nature. These proteins structurally transduce biochemical information in a manner analogous to how transistors control information flow in computing devices. Engineering challenges ranging from biological computing devices to molecular motors require such two-state switches, but designing these is an unsolved problem as it requires sculpting an energy landscape with two low-energy but structurally distinct conformations that can be modulated by external inputs. Here we describe a general design approach for creating “hinge” proteins that populate one distinct state in the absence of ligand and a second designed state in the presence of ligand. X-ray crystallography, electron microscopy, and double electron-electron resonance spectroscopy demonstrate that despite the significant structural differences, the two states are designed with atomic level accuracy. The kinetics and thermodynamics of effector binding can be finely tuned by modulating the free energy differences between the two states; when this difference becomes sufficiently small, we obtain bistable proteins that populate both states in the absence of effector, but collapse to a single state upon effector addition. Like the transistor, these switches now enable the design of a wide array of molecular information processing systems.

## Introduction

While many naturally occurring proteins adopt single folded states, conformational changes between distinct protein states are crucial to the functions of enzymes(*1, 2*), cell receptors(*3*), and molecular motors(*4*). The extent of these changes ranges from small rearrangements of secondary structure elements(*5, 6*) over domain movement(*7*) to fold-switching or metamorphic proteins(*8*) that adopt completely different structures. In many cases, these conformational changes are triggered by “input” stimuli such as binding of a target molecule, post-translational modification, or change in pH. These changes in conformation can in turn result in “output’’ actions such as enzyme activation, target binding, or oligomerization(*9*); protein conformational changes can thus couple a specific input to a specific output. The generation of proteins that can switch between two quite different structural states is a difficult challenge for computational protein design, which usually aims to optimize a single, very stable conformation to be the global minimum of the folding energy landscape(*10, 11*); design of proteins that can undergo controlled, major conformational changes requires reframing this paradigm towards optimizing for more than one minimum on the energy landscape, while simultaneously avoiding undesired off-target minima(*12*). Previously, multi-state design has been used to design proteins that undergo very subtle conformational changes(*13, 14*), cyclic peptides that switch conformations based on the presence of metal ions(*15*), and closely related sequences that fold into dramatically different conformations(*16*). Stimulus-responsive proteins have been designed to undergo conformational changes upon binding to a target peptide or protein(*17, 18*); however, while the “closed” unbound state of these “switch” proteins is a well-defined and fully structured conformation, the “open” bound state is a broad distribution of conformations. These proteins have found use as biosensors(*19, 20*), but the lack of a defined second state makes them not well suited for mechanical coupling in a molecular machine or discrete state based computing systems.

We set out to design proteins that can switch between two well-defined and fully structured conformations. To facilitate experimental characterization of the conformational change and to ensure compatibility with downstream applications, we imposed several additional requirements. First, the conformational change between the two states should be large, with some inter-residue distances changing by tens of angstroms between the two states. Second, the conformational change should not require global unfolding, which can be very slow. Third, neither of the two states should have substantial exposed patches of hydrophobic residues, which can compromise solubility. Fourth, the conformational change should be readily coupled to a range of inputs and outputs. Given that proteins are stabilized by hydrophobic cores, collectively achieving all of these properties in one protein system is challenging: protein conformations that differ considerably typically will have different sets of buried hydrophobic residues and require substantial structural rearrangements for interconversion.

We reasoned that these goals can collectively be achieved with a “hinge”-like design in which two rigid domains move relative to each other while remaining individually folded. The hinge amplifies small local structural and chemical changes to achieve large global changes while the chemical environment for most residues remains similar throughout the conformational change, avoiding the need for global unfolding. Provided that the two states of the hinge bury similar sets of hydrophobic residues, the amount of exposed hydrophobic surface area can be kept low in both states. Designing one of the resulting conformations to bind to a target effector couples the conformational equilibrium with target binding (Figure 1A). This design concept has precedent in nature; for example bacterial two-component systems utilize binding proteins that undergo hinging between two discrete conformations in response to ligand binding(*21*).

**Figure 1:**
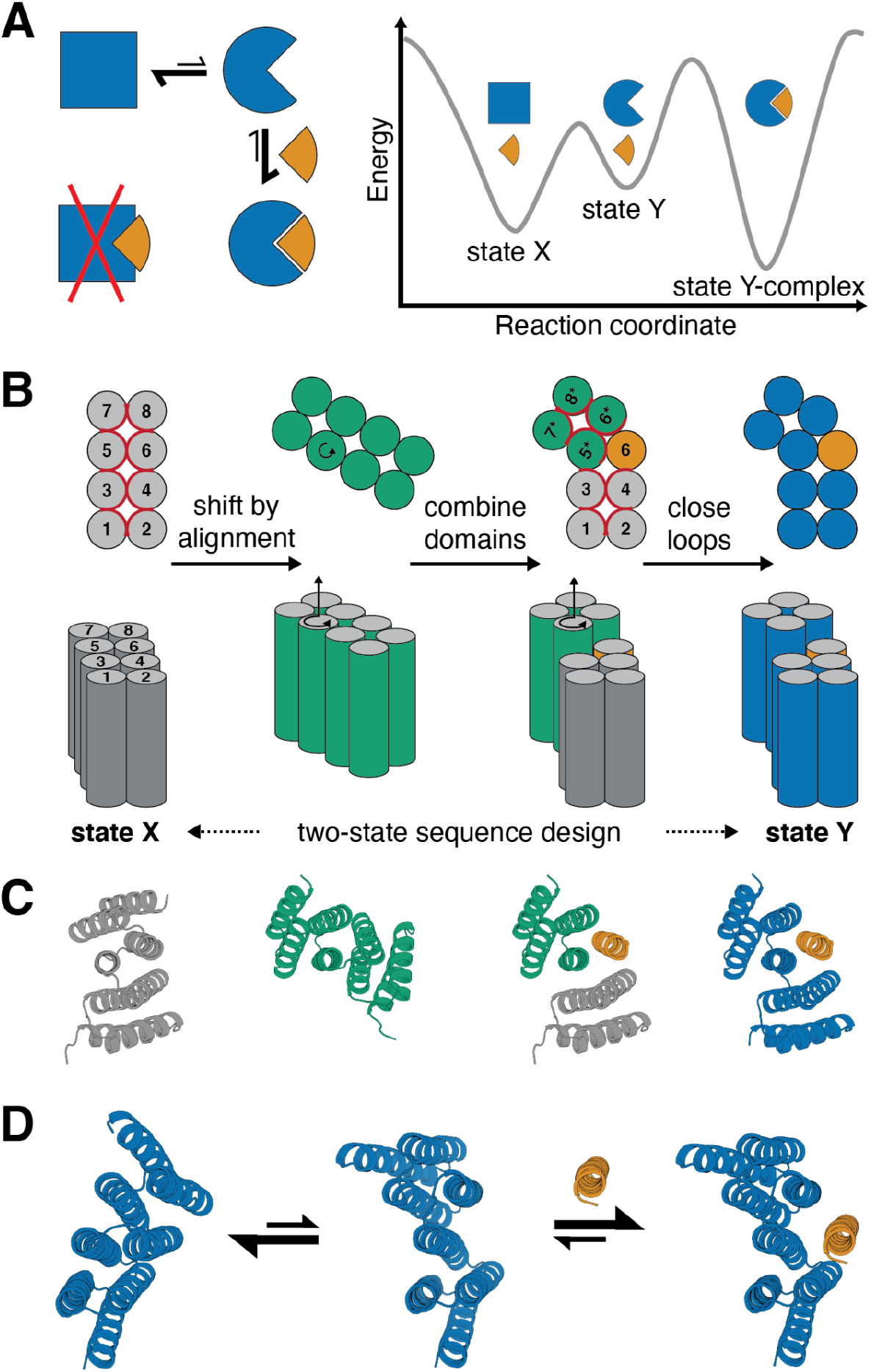
Strategy for designing proteins that can switch between different conformations. **A)** Left: reaction scheme for a protein (blue) that undergoes a conformational change and can bind an effector (orange) in one (circle) but not in the other conformational state (square). Right: Energy landscape for the system shown on the left. **B)** Schematic representation of the hinge design approach. Alpha-helices are represented as circles (top view, top) or cylinders (side view, bottom). From left to right: A previously designed repeat protein (gray) serves as the first conformation of the hinge. To generate the second conformation a copy of the repeat protein (green) is moved by shifted alignment along a pivot helix, causing a rotation (top and bottom, indicated by circular arrow) and a translation along the helix axis (bottom). The first 4 helices of the original protein form domain 1 of the hinge, the last 4 helices of the rotated copy form domain 2, and an additional helix is copied over from the original protein to serve as an effector peptide (orange) that can bind to this second conformation of the hinge. Both domains of the hinge are connected into one continuous chain (blue) using fragment-based loop closure, and a single amino acid sequence is designed to be compatible with both conformations. **C)** Design steps from B illustrated using cartoon representations of an exemplary design trajectory. **D)** Exemplary design models of a designed hinge protein in state X (left), state Y (center), and in state Y bound to an effector peptide (right). Hinge is shown in blue, peptide in orange.

## Design Method

To implement this two-state hinge design concept, we took advantage of designed helical repeat proteins (DHRs, (*22*); Figure 1B,C left) or DHR-based junction proteins(*23*). The backbone conformation of the DHR serves as the first conformational state of our hinge protein (“state X”). To generate a second conformation, a copy of the parent protein is rotated around a “pivot helix” by aligning the copy to the original DHR shifted by N residues, where −7 < N < 7 (Figure 1B,C). A new backbone conformation is then created by combining the first half of the original protein (domain 1), the second half of the copy (domain 2), and either the helix following the pivot helix from the original protein or the helix preceding the pivot helix from the rotated copy (“peptide”). Backbone arrangements with large backbone clashes, as evaluated by the Lennard-Jones potential in Rosetta(*24*), are discarded. Rosetta FastDesign with backbone movement(*25, 26*) is used to re-design the interface between the three parts. Using fragment-based loop closure(*22, 27, 28*), domains 1 and 2 are connected into a single chain that serves as the second conformational state of the hinge protein (“state Y”, Figure 1B,C right). The loop between domains 1 and 2 is rebuilt in state X, yielding pairs of state X and state Y backbones with matching loop lengths and secondary structures. Using a combination of Rosetta two-state design (see methods section for details) and proteinMPNN(*29*) with linked residue identities, a single amino acid sequence is generated that is compatible with the state X hinge as well as with the state Y hinge-peptide complex. AlphaFold2 (AF2)(*30*) with initial guess(*31*) can then be used to predict the structure of the hinge with and without the effector peptide, allowing for the selection of designs that are predicted in the correct state X in absence of the peptide and in the correct state Y complex in presence of the peptide. The designs are additionally filtered based on the Rosetta energies and spatial aggregation propensity (SAP)(*32*) for state X and for state Y with and without peptide. To favor designs that are predominantly in the closed state in absence of the peptide (Figure 1A,D), designs are selected only if state X has lower energy than state Y in absence of the peptide, and if the state Y complex has lower energy than state X plus the peptide, spatially separated. Designs are also filtered on standard interface design metrics for the bound conformation (see methods for details on filtering)(*33*).

## Hinges bind effector peptides with sub-nM to low μM affinities

We used our hinge design approach to generate hinge-peptide pairs that cover a large structural space with diverse changes in terminal angle and globularity (Figures 1D, 2A, S1, S3). We experimentally tested multiple rounds of designs, using both DHRs(*22*) and helical junctions(*23*) as input scaffolds, and improving individual steps of the design pipeline between iterations (see Methods). To facilitate expression and improve solubility, the peptides were fused to superfolder GFP (sfGFP) via a flexible linker. Hinges and sfGFP-peptide fusions were expressed in *Escherichia coli* (E. coli) and purified using immobilized metal affinity chromatography (IMAC) followed by size exclusion chromatography (SEC). For hinge designs that were predominantly monomeric by SEC at high expression levels, we evaluated peptide binding by running hinge, the corresponding peptide-sfGFP fusion, and a mixture of both on SEC (Figures S2, S3). Designs that showed significant peak shifts of the mixture compared to individual components, indicating hinge-peptide binding, were selected for further characterization using a fluorescence polarization (FP) assay in which a chemically synthesized 5-carboxy-tetramethylrhodamine (TAMRA)-labeled peptide was incubated with different concentrations of purified hinge protein (Figures 2B, S4). The measured polarization is proportional to the fraction of bound peptide. Hinge-peptide binding affinities obtained from titration experiments with constant peptide concentration and varying hinge concentrations ranged from 1 nM to the low μM range (Figures 2B, S4). To circumvent the bottleneck of finding soluble peptide sequences we sought to design hinges that bind to a given target peptide. Starting from hinge cs201, we used a modified version of our design pipeline to redesign the hinge to bind the peptide from designs cs074 or cs221, respectively. This one-sided two-state design approach yielded hinge designs that showed strong binding to their new target peptide while showing no or only weak off-target binding (Figure S5).

**Figure 2:**
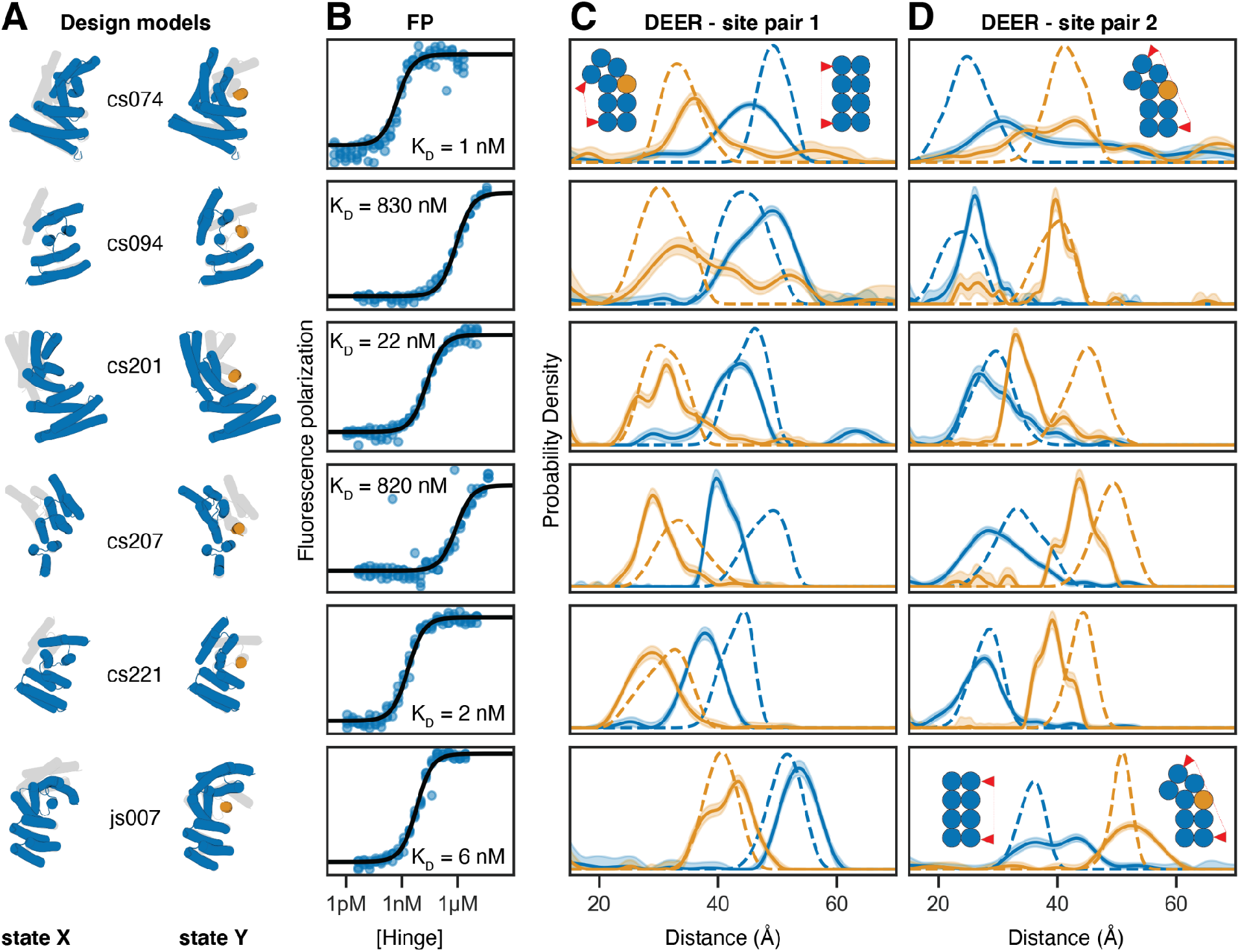
Experimental validation of peptide-binding hinges. **A)** Design models of hinges (blue) and peptides (orange) in state X (left model) and state Y bound to the peptide (right model). Gray shades behind models in state X and Y indicate the corresponding states Y and X, respectively. **B)** Fluorescence Polarization (FP) titrations with a constant concentration of TAMRA-labeled peptide (0.1 nM for cs074 and cs221; 0.5 nM for cs201; 1 nM for cs094, cs207, and js007) and varying hinge concentrations. Circles represent data points from four independent measurements, lines are fits of standard binding isotherms to all data points, dissociation constants (K_D_) are obtained from those fits. **C,D)** Distance distributions between spin labels covalently attached to cysteine side chains. Solid lines are obtained from DEER experiments without (blue) or with (orange) an excess of peptide, shaded areas are 95% confidence intervals, and dashed lines are simulated based on the design models for state X (blue) or the state Y complex (orange). For each hinge two different label site pairs were tested, one in which the distance was expected to decrease with peptide binding (C) and one in which the distance was expected to increase upon peptide binding (D). Chemically synthesized peptides were used for all measurements except for cs074 site pair 1, for which sfGFP-peptide fusion was used.

## Effector binding modulates the hinge conformational equilibrium

To characterize the conformational equilibrium of the designed hinges, we introduced two surface cysteine residues into the hinge protein and covalently labeled them with the nitroxide spin label MTSL(*34*). We then used double electron-electron resonance spectroscopy (DEER) to determine distance distributions between the two spin labels and compared these to simulated(*35*) distance distributions based on the state X and state Y design models. This experiment was performed on two different labeling site pairs for each design: one pair where the distance is predicted to decrease in the presence of peptide (Figures 2C, S4) and the other where it is predicted to increase (Figures 2D, S4). In the absence of the peptide, the observed distance distributions closely matched the state X simulations. In all cases the distances between the two pairs of probes shifted upon addition of peptide to better match the state Y simulations, suggesting that addition of effector peptide causes the conformational equilibrium to shift towards state Y as designed. For example, cs074 (site pair 1) showed a clear peak between 40 and 50 Å in absence of the peptide, and a peak between 30 and 40 Å in presence of the peptide, and both peaks agree well with the corresponding simulations (Figure 2C, top row). In a control experiment using the static parent DHR protein of design cs074, the distance distributions with and without peptide were identical and matched both the simulation for the parent design model, which closely resembles state X, and the experimental distance distribution for state X of cs074 (Figure S4). Design cs094 (site pair 1) showed the expected state X peak around 50 Å in absence of the peptide. In presence of peptide this design showed a new peak around 30 Å matching the state Y simulation, but still showed a residual state X peak around 50 Å, indicating incomplete binding either due to weak binding affinity or to insufficient peptide concentration. For all hinge designs tested using DEER, the observed conformational changes were consistent with the designed state X and Y models.

We solved crystal structures for two designs, cs207 and cs074. For design cs207, crystals were obtained from two separate crystallization screens: one screen for the hinge alone (Figure 3A), and one screen for the hinge in complex with the target peptide (Figure 3B). In the absence of peptide the experimental structure agrees well with the state X design model, with an RMSD (root mean square deviation) of 1.03 Å between model and experimental structure at a resolution of 2.50 Å. The structure of the hinge-peptide complex agrees well with the state Y design model with an RMSD of 0.69 Å between model and experimental structure at a resolution of 2.66 Å. For comparison, the RMSD between design models for states X and Y of cs207 is 4.59 Å. The electron densities for the side chains involved in the interdomain interface in state X and in the hinge-peptide interface in state Y closely match the design models, providing high-resolution validation of our two-state design procedure (Figure 3A-C, right). The two crystal structures of cs207 with and without peptide, combined with our DEER distance distributions, corroborate the fundamental hinge design concept: In the absence of the peptide the hinge is predominantly in state X, and addition of the peptide shifts the equilibrium towards the state Y-peptide complex. For another design, cs074, the crystal structure of the hinge-peptide complex (Figure 3C) agrees well with the corresponding state Y design model (RMSD 1.46 Å to original Rosetta model, 0.67 Å to the closest AF2 prediction, at a resolution of 2.75 Å).

**Figure 3:**
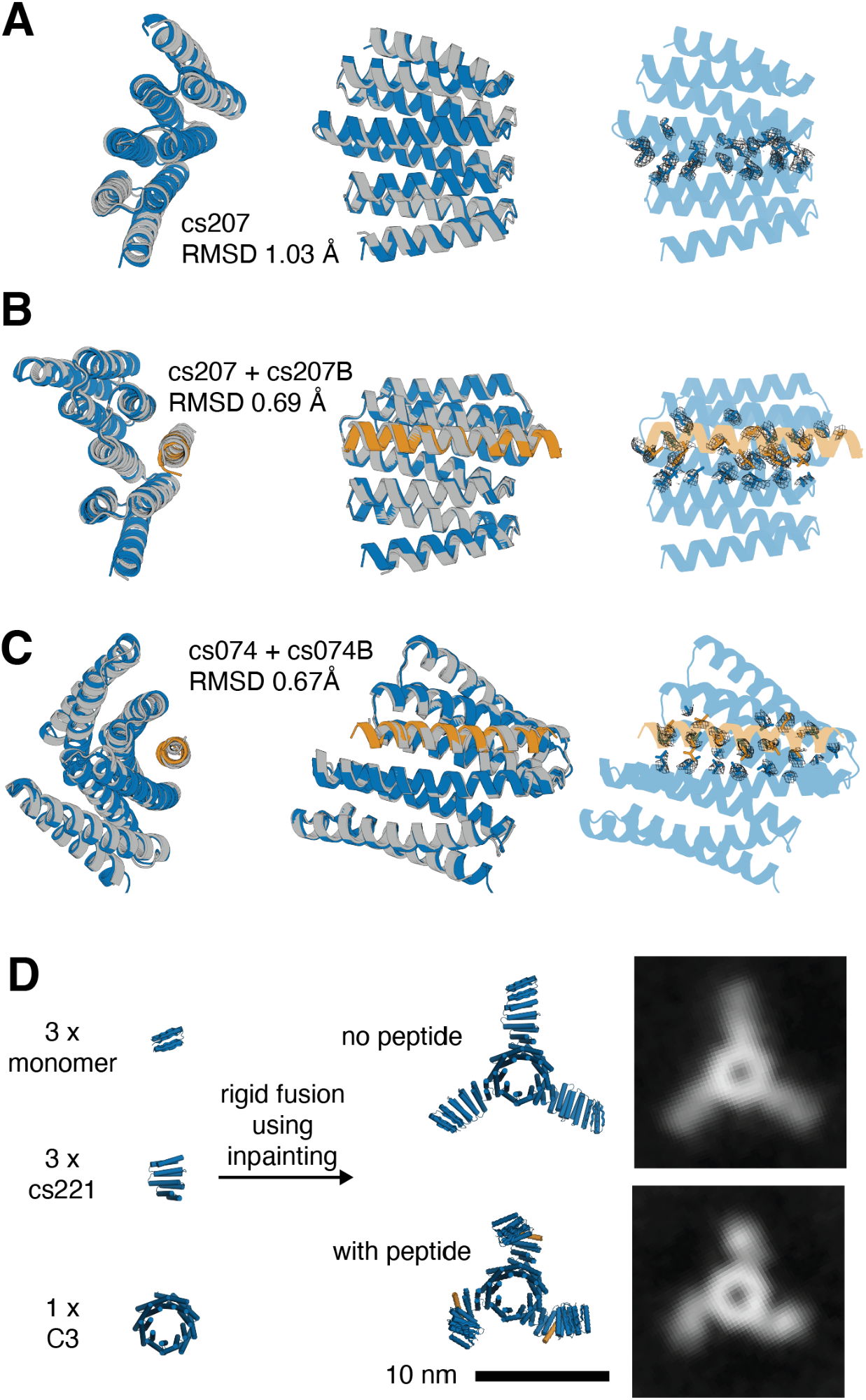
Structural validation of the conformational change in peptide-binding hinges. **A,B,C)** Design models (hinge in blue, peptide in orange) overlaid with crystal structures (gray) in top view (left) and side view (center) and design models with electron density for selected side chains (right). **A)** Design model of hinge cs207 in state X overlaid with crystal structure of hinge cs207 crystallized without peptide. Right panel highlights electron densities for side chains in the interface between the two hinge domains. **B)** Design model of hinge cs207 in state Y overlaid with crystal structure of hinge cs207 crystallized with peptide cs207B. Right panel highlights electron densities for the same side chains as A; these side chains now contribute to the hinge-peptide interface. **C)** Design model of hinge cs074 in state Y overlaid with crystal structure of hinge cs074 crystallized with peptide cs207B. Right panel highlights electron densities for side chains in the interface between hinge and peptide. **D)** Left: Components for design of a C3-symmetric homotrimer with three cs221 hinge arms. Center: Design model of the hinge-armed trimer in state X (top) and in state Y (bottom). Right: nsEM class averages of the trimer in absence of peptide (top) and in presence (bottom) of peptide cs221B.

To test whether our hinges can be incorporated as components of more complex protein assemblies without affecting their ability to undergo conformational changes, we designed a fully structured C3-symmetric protein with three hinge arms (Figure 3D). We used inpainting(*36*) with RoseTTAFold(*37*) to rigidly connect one end of hinge cs221 to a previously validated homotrimer(*38*) and the other end of the hinge to a previously validated monomeric protein(*39*). Negative-stain electron microscopy (nsEM) with reference-free class averaging shows straight arms in absence of peptide and bent arms in presence of peptide cs221B, corroborating the designed conformational change (Figures 3D, S6).

A critical feature of two-state switches in biology and technology is the coupling between the state control mechanism and the populations of the two states. To quantitatively investigate the thermodynamics and kinetics of the effector induced switching between the two states of our designed hinges, we used Förster resonance energy transfer (FRET). To increase both the absolute distance from N- to C-terminus and the change in termini distance between the two conformational states, we took advantage of the extensibility of repeat proteins and extended hinges cs074, cs221, and cs201 by 1-2 helices on their N and C termini, yielding cs074F, cs221F, and cs201F, respectively (Figure 4A, first column). Single cysteines were introduced in helical regions near the termini of the extended hinges and stochastically labeled with an equal mixture of donor and acceptor dyes. For hinges cs074F and cs221F the distance between the label sites is above the R0 of the dye pair in state X and below R0 in state Y, and hence, acceptor emission upon donor excitation increases upon addition of the corresponding peptides cs074B and cs221B, respectively (Figure 4A, second column). We used labeled, extended DHR82, the parent protein for cs074F, as a static control, and observed fluorescence spectra comparable to cs074F but no change in fluorescence upon addition of the peptide (Figure S7). For cs201F, the dye distance is above R_0_ in state X and below R_0_ in state Y, and donor emission decreases upon addition of peptide cs201B (Figure 4A, second column). To test specificity of our hinge-peptide pairs, we performed pairwise titrations of all three labeled hinges at 2 nM with all three target peptides at varying concentrations. The on-target titrations had sigmoidal transitions that can be fitted with standard binding isotherms (Figures 4A, third column; S7), while the off-target titrations for cs201F and cs221F show flat lines, indicating no conformational change of these hinges upon addition of off-target peptides at μM concentrations. cs074F showed weak off-target binding that was three orders of magnitude weaker for cs201B and two orders of magnitude weaker for cs221B compared to the on-target interaction for cs074B. cs201F and cs221F are thus orthogonal in the nM and μM range, and the set of cs201F, cs221F, and cs074F is orthogonal over two orders of magnitude of effector concentration.

**Figure 4:**
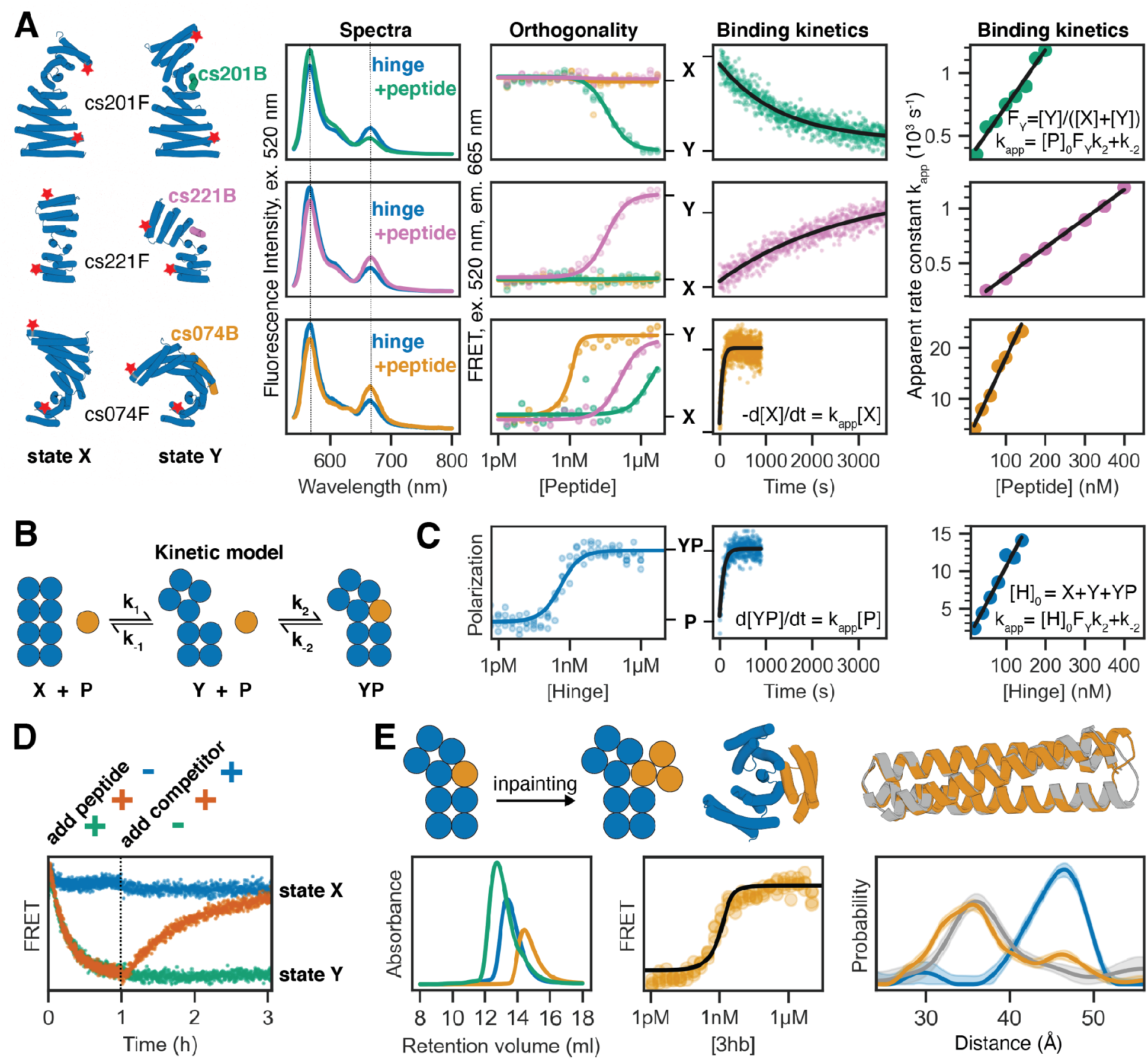
Quantitative analysis of conformational changes in designed hinge proteins. **A)** FRETbased characterization of three extended hinges. From left to right: cylindrical representation of extended hinges (blue) and their corresponding target peptides (green: cs201B, pink: cs221B, orange: cs074B) with red stars indicating attachment sites for fluorescent dyes; fluorescence spectra (excitation at 520 nm) of labeled hinge without (blue) or with (green/pink/orange) target peptide; FRET-based binding titrations (excitation 520 nm, emission 665 nm) at 2 nM labeled hinge and varying peptide concentrations fitted with standard binding isotherms (solid lines); time course after mixing 2 nM (cs201F, cs074F) or 5 nM (cs221F) labeled hinge and 100 nM peptide fitted with a single-exponential equation (black line); apparent rate constants obtained from single-exponential kinetic fits plotted against absolute peptide concentrations (circles) and fitted with a linear equation (black line). Dotted lines in spectra indicate acceptor and donor emission peaks. **B)** Kinetic model describing the coupling of the conformational equilibrium to the binding equilibrium. X and Y: hinge in state X and Y, respectively; P: peptide; YP: peptide bound to hinge in state Y. k_1_, k_-1_, k_2_, and k_-2_ are the microscopic rate constants. **C)** FP characterization of unlabeled extended hinge cs074F. From left to right: binding titration at 0.1 nM TAMRA-labeled peptide and varying hinge concentrations; time course after mixing 2 nM TAMRA-labeled peptide and 100 nM hinge fitted with a single-exponential equation (black line); apparent rate constants obtained from single-exponential kinetic fits plotted against absolute hinge concentrations (circles) and fitted with a linear equation (black line). **D)** FRET-based reversibility experiment using the labeled extended hinge cs201F introduced in C). Hinge concentration is 30 nM for all traces; 1 μM peptide is added at t=0 (green/orange), 3 μM unlabeled competitor hinge is added after 1 h (blue/orange). **E)** Top from left to right: schematic representation of the inpainting procedure that adds two helices to the peptide cs074B yielding a three-helix bundle (3hb); cylindrical representation of 3hb_05(orange) bound to hinge cs074 (blue); overlay of design model (orange) and crystal structure (gray) of 3hb_05. Bottom from left to right: SEC traces for hinge cs074 (blue), 3hb_05 (orange), and a mixture of both (green); FRET-based titration of 2 nM extended labeled hinge cs074F and varying concentrations of 3hb_05 fitted with a standard binding isotherm (back line); Distance distributions obtained from DEER experiments as described in Figure 2 (blue: cs074, gray: cs074 + peptide cs074B, orange: cs074 + 3hb_05).

Association kinetics for the on-target interactions measured using constant concentrations of labeled hinge and varying excess concentrations of peptide are well fit by single exponentials (Figures 4A, fourth column; S8). The apparent rate constants increase linearly with increasing peptide concentration, exhibiting standard pseudo-first order kinetics for bimolecular reactions (Figures 4A, fifth column; S8). We analyze these data using a model comprising the three states (X, Y, Y+peptide) and four rate constants (Figure 4B). The kinetic measurements using the FRET system follow the decrease in state X over time (d[X]/dt) upon the addition of peptide. The observed pseudo-first order behavior (Figure 4A, fifth column) indicates that the conformational change happens on a timescale that is faster than that of the observed binding and can be treated as a fast pre-equilibrium (Supplementary Note 1). The slopes of the linear pseudo-first order fits (k_on_) can thus be interpreted as the product of the microscopic association rate k_2_ and the fractional population of state Y in absence of the peptide (F_Y_ = [Y]/([X]+[Y]), see Supplementary Note 1). FP based titrations and kinetic characterization using the unlabeled extended hinge cs074F in excess over the TAMRA-labeled peptide cs074B agree well with the corresponding FRET experiments, further supporting the preequilibrium model (Figures 4C, S8). FP kinetics experiments for other hinge designs also follow pseudo-first order behavior with k_on_ values ranging from 2.5×10^3^ M^-1^s^-1^ to 7.8×10^4^ M^-1^s^-1^ (Figures S4, S9). To study the reversibility of hinge conformational changes, we started with 30 nM of FRET-labeled hinge cs201F (Figure 4D), added 200 nM peptide to drive the conformational change, and then added excess unlabeled hinge cs201 to compete away the peptide. The FRET signal decreased upon addition of the peptide, consistent with conformational change from state X to state Y, and then returned to nearly the original level upon addition of unlabeled hinge, indicating that the hinge conformational change is fully reversible.

To explore whether peptide-responsive hinges could be turned into protein-responsive hinges, we used inpainting with RoseTTAFold to add two additional helices to a validated effector peptide, resulting in fully structured 3-helix bundles (3hb). For nine of our validated hinges, we designed and experimentally characterized these effector proteins using SEC (Figures 4E, S10, S11). Hinge-3hb binding was tested qualitatively by SEC and, for hinges which had a corresponding FRET construct, quantitatively with the FRET-labeled variant, and DEER was used in addition to FRET to confirm that 3hb binding caused the same conformational change as effector peptide binding (Figures 4E, bottom; S10). The affinity of 3hb05 to cs074F was similar to the affinity observed for the original peptide cs074B (Figure 4E), whereas 3hb21 bound its target hinge cs221F significantly tighter than the original peptide cs221B (Figure S12). The 3hb approach was able to rescue designs for which the peptide alone or the hinge-peptide complex had shown the tendency to form higher-order oligomers (Figure S11). For two designs, 3hb05 and 3hb12, we obtained crystal structures that agreed well with the design models, indicating that the three-helix bundles are fully structured in isolation (Figures 4E top right, S13).

## The conformational pre-equilibrium controls effector binding

To test the effect of the conformational pre-equilibrium on effector binding, we introduced disulfide “staples” that lock the hinge in one conformation. We used a 6D hashing approach(*40*) to identify pairs of cysteine positions that would be compatible with disulfide formation in one state but not the other, and used AF2 prediction as a computational filter, selecting only sequences for which cysteine side chains were predicted to have side chain distances and relative orientations compatible with disulfide formation in the intended structural state. Using FP we analyzed peptide binding to stapled versions of hinge cs221 (Figure 5A,B). The variant that forms a disulfide bond in state X (“locked X”) showed only weak residual binding, likely due to a small fraction of hinges not forming the disulfide (Figure 5A). Upon addition of the reducing agent dithiothreitol (DTT) to break the disulfide, peptide binding was fully restored, making this hinge variant a red/ox dependent peptide binder that binds the effector peptide under reducing but not under oxidizing conditions. The variant that forms a disulfide bond in state Y (“locked Y”) showed fast peptide binding following pseudo-first order kinetics with an observed association rate that was 200-fold higher than for the original hinge without disulfides (Figures 5B, S14). Using the pre-equilibrium model described above, the observed association rates provide an estimate of the fraction of hinge that is in state Y in absence of the peptide: a 200-fold higher observed on rate for the locked Y variant indicates a 200-fold higher fraction of hinge in state Y compared to the original hinge. Assuming that the locked Y variant is 100% in state Y and assuming that the microscopic rate constant k_2_ is identical for the locked Y hinge and state Y of the original hinge, this would indicate that the original hinge is 99.5% in state X and 0.5% in state Y at equilibrium.

**Figure 5:**
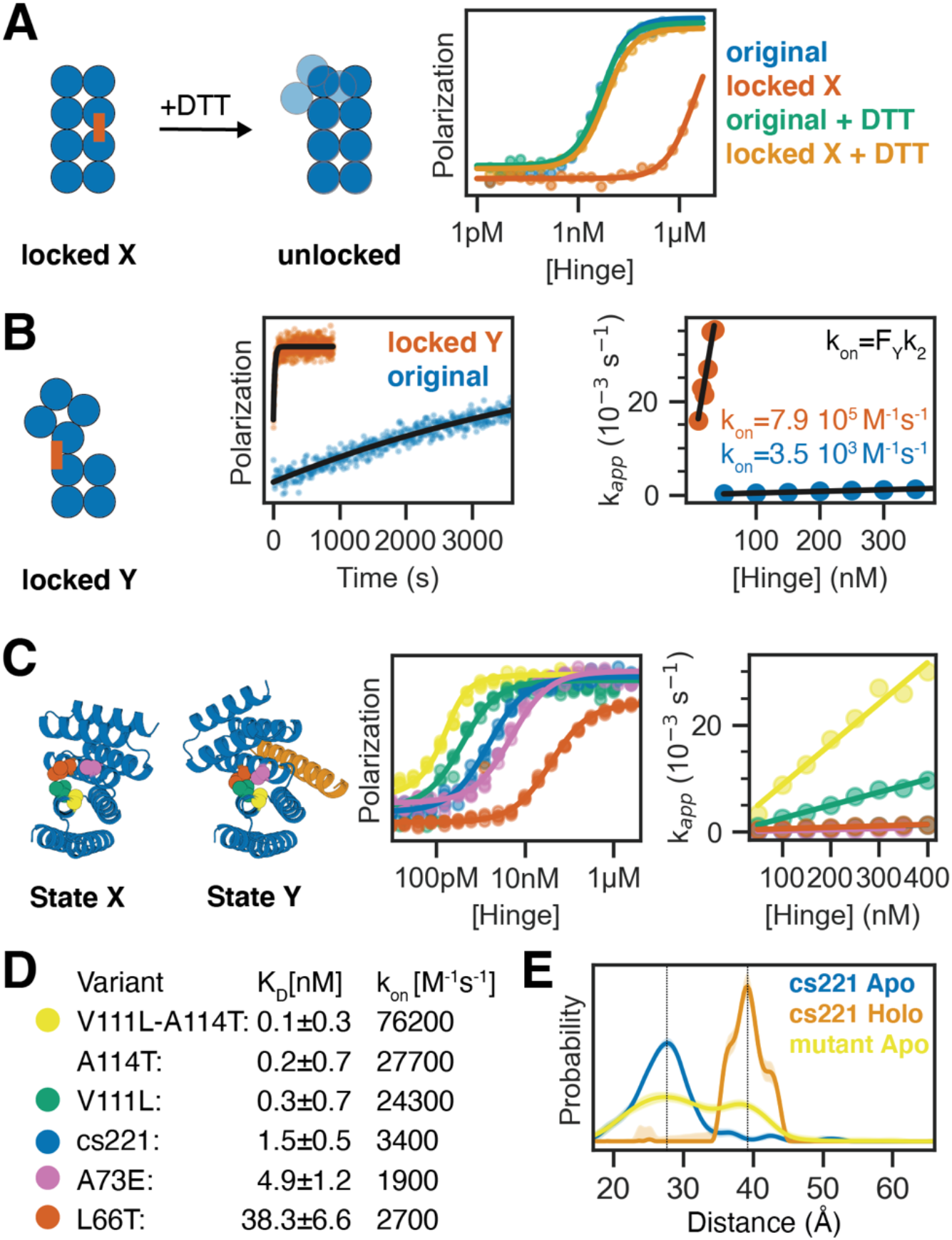
Controlling the conformational pre-equilibrium affects peptide binding. **A)** Left: Schematic representation of a hinge containing two cysteine residues that can form a disulfide bond in state X but not in state Y, effectively locking the hinge in state X under oxidizing conditions. Upon addition of reducing agent DTT the disulfide bond is broken and the conformational equilibrium is restored. Right: FP-based titration of 1 nM TAMRA-labeled peptide and a hinge with state X disulfide (red, orange) or the parent hinge without cysteines (blue, green) under oxidizing (blue, red) or reducing (green, orange) conditions. **B)** From left to right: schematic representation of a hinge that is disulfide-locked in state Y; time course after mixing 2 nM TAMRA-labeled peptide and 50 nM locked hinge (red) or original hinge without cysteines (blue) fitted with a single-exponential equation (black line); apparent rate constants obtained from single-exponential kinetic fits plotted against absolute hinge concentrations (circles) and fitted with a linear equation (black line). **C)** Tuning the pre-equilibrium with point mutations. Left: Cartoon representation of hinge cs221 highlighting positions of point mutations. Center: FP-based titration of 0.1 nM (yellow, green, blue) or 1 nM (pink, red) TAMRA-labeled peptide cs221B and varying concentrations of hinge variants containing one or two point mutations. Right: Apparent rate constants obtained from single-exponential kinetic fits plotted against absolute hinge concentrations (circles) and fitted with a linear equation (black line). **D)** Dissociation constants (K_D_) and observed binding rate constants (k_on_) for the hinge variants shown in C. **E)** DEER distance distribution for the double mutant cs221-V111L-A114T in absence of peptide (yellow) in comparison to the original cs221 with (orange) and without (blue) peptide. Gray lines serve as guide to the eye indicating state X and state Y distances.

Having established the edge cases of locked state X and locked state Y, we sought to tune the pre-equilibrium by introducing single point mutations expected to specifically stabilize one state over the other while not directly affecting the peptide-binding interface. We used proteinMPNN to generate consensus sequences(*41*) for each state and identified non-interface positions with distinct residue preferences that were different between both states (Figure 5C). We experimentally tested individual protein variants carrying mutations expected to stabilize one state over the other without disrupting either conformation, as evaluated by AF2 predictions. Several mutations expected to stabilize state X led to weaker peptide binding, and mutations expected to stabilize state Y led to stronger binding (Figures 5C,D, S14). Mutations expected to stabilize state X had little effect on k_on_, suggesting that these mutations primarily caused destabilization of the state Y-peptide complex. Mutations that stabilized state Y, on the other hand, strongly affected k_on_, indicating that they effectively shift the pre-equilibrium between both states (Figures 5C,D, S15).

The double mutant cs221_V111L_A114T has a 22-fold higher on rate than the original cs221, suggesting the occupancy of state Y in cs221_V111 L_A114T is 22x higher in the absence of peptide. Distance distributions obtained from DEER measurements on site pair 2 of the double mutant cs221_V111L_A114T in absence of the peptide indeed showed an additional peak at a distance closely matching state Y (Figures 5E, S16). DEER measurements on site pair 1 of the double mutant showed a broader distribution with occupancy in the region corresponding to state Y (Figures 5E, S16). Measurements in the presence of the peptide were virtually indistinguishable from the original cs221 (Figure S16). The double mutant thus populates two distinct states in the absence of the effector, and collapses to one state upon effector addition (Figures 5E, S16). The observation of a significant state Y population at equilibrium in the absence of the peptide as predicted based on the kinetic measurements further corroborates that the mutations affect the conformational pre-equilibrium, and provides strong support for our quantitative two-state model of the kinetics and thermodynamics of the designed hinge-effector systems.

## Conclusion

Taken together, our structural, kinetic and thermodynamic data strongly suggest that our hinge design method generates proteins that populate two well-defined and structured conformational states. The crystal structures, nsEM class averages, and one-dimensional distance distributions obtained from DEER measurements suggest that the designed hinges preferentially populate the two designed states, rather than adopting a heterogenous mixture of structures. The DEER and FRET experiments show that a peptide or protein effector can drive the conformational equilibrium from state X to state Y. The kinetics of the conformational change measured by FRET and the kinetics of peptide binding measured by FP agree well, corroborating the two-state model. DEER measurements on a hinge variant that populates both states in absence of the peptide further confirm the absence of additional states and agree well with population estimates based on association kinetics. To our knowledge these are the first designed proteins that can reversibly switch between two substantially different structured conformations. Our design strategy for creating protein two-state systems without exposing substantial hydrophobic surface area in either state should be quite broadly applicable.

Like transistors in electronic circuits, our designed two-state switches can now be coupled to external outputs and inputs to create sensing devices and incorporated into larger protein systems to address a wide range of outstanding design challenges. Hinges containing a disulfide that locks them in state X couple the input “red/ox state” to the output “target binding,” where the target can be a peptide or a protein, and our FRET-labeled hinges couple the input “target binding” to the output “FRET signal.” Our approach can be readily extended to have state switching driven by naturally occurring rather than designed peptides; hinge-like proteins have recently been designed to target peptides such as glucagon, secretin, or neuropeptide Y(*42*), enabling new routes to sensing and detection. Stimuli-responsive protein assemblies that change shape or oligomeric state in the presence of an effector can now be built by incorporating the hinges as modular building blocks. Installing enzymatic sites in hinges such that substrate binding favors one state and product release favors the other state should enable fuel-driven conformational cycling, a crucial step towards the *de novo* design of molecular motors. More generally, the ability to design two-state systems, and the designed two-state switches presented here, should enable protein design to go beyond static structures to more complex multistate assemblies and machines.

## Acknowledgements

We thank Basile I. M. Wicky, Lukas F. Milles, and Danny D. Sahtoe for helpful discussions and technical support, Alexis Courbet for inspiring discussions, Annika Philomin and Andrew Borst for EM support, and Kandise VanWormer and Luki Goldschmidt for technical support.

We want to thank the Advanced Light Source (ALS) beamline 8.2.2/8.2.1 at Lawrence Berkeley National Laboratory for X-ray crystallography data collection. The Berkeley Center for Structural Biology is supported in part by the National Institutes of Health (NIH), National Institute of General Medical Sciences, and the Howard Hughes Medical Institute. The ALS is supported by the Director, Office of Science, Office of Basic Energy Sciences and US Department of Energy (DOE) (DE-AC02-05CH11231).

## Funding

This work was supported by a Human Frontier Science Program Long Term Fellowship (LT000880/2019, F.P.), the Open Philanthropy Project Improving Protein Design Fund (P.J.Y.L., C.W.D., H.N., D.B.), an NSF Graduate Research Fellowship (DGE-2140004; P.J.Y.L), NERSC award BER-ERCAP0022018 (P.J.Y.L, D.B.), the Audacious Project at the Institute for Protein Design (A.B., A.P., A.I., M.L., R.K.B., S.R.G., A.K., D.B.), a gift from Microsoft (D.J., J.D., D.B.), a grant from DARPA supporting the Harnessing Enzymatic Activity for Lifesaving Remedies (HEALR) program (HR001120S0052 contract HR0011-21-2-0012, X.L., A.K.B., D.B.), the Defense Threat Reduction Agency (DTRA) grant # HDTRA1-19-1-0003 (P.M.L.), NSF Award #2006864 (J.N.), and the Howard Hughes Medical Institute (D.B.). DEER measurements were supported by R01 GM125753 (to S.S.). The spectrometer used was funded by NIH grant S10 OD021557 (S.S.).

## Author contributions

F.P. developed the hinge design concept. F.P. and P.J.Y.L. developed the computational hinge design pipeline. F.P. and P.J.Y.L. designed, screened, and characterized most hinges with help from C.D. and A.P.. M.H.T. designed, performed and analyzed DEER experiments. S.S. analyzed DEER data and supervised research. C.D. developed the one-sided two-state design protocol and designed and tested hinges with swapped targets. A.B. designed and characterized 3-helix bundles with support from D.C.J.. A.I. designed and characterized the hinge-armed trimer with support from A.B.. A.I. performed electron microscopy and image processing with support from A.P.. J.D. provided conceptual support for two-state sequence design. X.L., P.M.L., M.L., and R.K.B. synthesized and purified peptides. X.L., M.L., and R.K.B. performed LC-MS validation of proteins and peptides. S.R.G. performed additional protein purification. H.N., A.K., B.S., and A.K.B. determined crystal structures. A.F.D. and B.V. contributed conceptual support. D.B. and J.N. supervised research. F.P., P.J.Y.L., and D.B. wrote the manuscript. M.H.T. contributed to the manuscript. All authors read and commented on the manuscript.

## Competing interests

A provisional patent application will be filed prior to publication, listing F.P., P.J.Y.L., M.H.T., A.B., C.D., A.F.D., A.P., A.I., B.V., S.S., and D.B. as inventors or contributors.

## Data and materials availability

All data are available in the main text or as supplementary materials. Design models and scripts are available at https://github.com/proleu/hinge_paper. Crystallographic datasets have been deposited in the Protein Data Bank (PDB) (accession codes 8FIH, 8FVT, 8FIT, 8FIN and 8FIQ).

## Notes

### Competing Interest Statement

A provisional patent application will be filed by the University of Washington, listing F.P., P.J.Y.L., M.H.T., A.B., C.D., A.F.D., A.P., A.I., B.V., S.S., and D.B. as inventors or contributors.

