## Supplementary materials for "Design of stimulus-responsive two-state hinge proteins"

**The PDF file includes:**

Materials and Methods
Supplementary Note S1
Figs. S1 to S18
Tables S1 to S2
References 44 - 60

### Methods

**Generating and pairing hinge conformational states**

We used curated libraries of DHRs as inputs for generation of hinge conformations. The backbone conformation of a given DHR serves as a template for the first conformational state (state X) of the hinge. The alignment-based alternative state generation protocol was implemented using custom PyRosetta functions. To discard backbone arrangements with significant clashes, we mutate the entire backbone to glycine and score the resulting pose with only the Rosetta “fa_rep” scoreterm(*24*), set to a weight of 1.0, with the “beta_nov16” scorefunction. At this stage, the helical peptide was sometimes extended using a similar alignment/shifting strategy to increase the size of the interface in state Y. We used PyRosetta FastDesign with backbone and jump movement to further improve the backbone and sequence around the tripartite interface between the first hinge domain, the peptide, and the second hinge domain for scoring purposes. At this stage, we discarded any designs where either of the domains or the peptide didn’t form good contacts with the other two chains using interface metrics(*33*). We called the blueprint builder(*43*) in PyRosetta to rebuild the loop region between the hinge domains in state X, based on a secondary structure template of the hinge in state Y.

**Two-state sequence design**

Initially, we tried many different multi-state design (MSD) algorithms in Rosetta. We first tried an approach where we would iterate between conformational states while performing single-state design (SSD) for each state individually while ramping a custom sequence convergence scoreterm between iterations(*44*). We found this method tended to decorate the surface with hydrophobics in positions that had ambiguous residue-level preferences between the conformational states, so we explicitly penalized excessive surface hydrophobics using constraints that calculated spatial aggregation propensity (SAP)(*45*) on the fly during design. We also used a quasisymmetric multistate design approach in PyRosetta, performing design on both states simultaneously while forcing the packer to consider the chemical context of residue positions linked across the states(*46*). This method seemed to have fewer pathologies in terms of positional sequence selection but scaled poorly in terms of computational performance, so we chose not to use it for large-scale sequence design tasks. Ultimately, we extensively used FastDesign with a version of the annealer originally intended and optimized for multi-conformation, sequence-symmetric design(*47*), since it was the easiest to use and scaled well computationally while being easily tunable to avoid the pathologies of the iterative approach. Once we had sampled sequences and backbones with Rosetta we optionally refined the sequences with proteinMPNN(*29*) multistate design (MPNN-MSD). Using a feature intended for homooligomer symmetry(*48*) we tied corresponding residue positions probabilities together across chains and used MPNN to sample up to 96 sequences per pair of backbones. We then could use AF2 initial guess (AF2-IG)(*31*) to predict the structure of the effector-bound complex (state Y) by threading the MPNN-MSD sequences back onto the backbones using mean predicted Local Distance Difference Test scores (pLDDT), RMSD to reference design model, and mean off-diagonal Predicted Aligned Error matrix (PAE interaction) cutoffs of 93, 1.5, and 5, respectively for AF2. Designs that passed these criteria could be predicted again by AF2(*30*, *49*) to check if they folded to the correct closed position (state X) absent the effector sequence. We observed that sequences designed with MPNN-MSD had much better computational success rates and overall metrics. After two-state design, most hinges were soluble and had significant monomer populations, however many of the effector peptides turned out to be insoluble or formed stable homooligomers. In some cases effector peptides were improved by simple redesign of the peptide surface residues away from the hinge interface, or by truncation of the peptide.

**Computational filtering**

Concerned with the possibility that hinges designed with this process would randomly oscillate between closed and open conformations in the absence of the effector, we tried to implement additional filters to select only the designs that would have our intended behavior for testing. We chose designs where the hinge sequence scored more favorably in Rosetta(*24*) in the closed conformation relative to the open conformation when the peptide was absent, but scored more favorably in the open conformation with the peptide bound in comparison to the sum of the scores of the closed conformation and the peptide alone. Similarly, we required that the solvent-exposed hydrophobicity, (measured by SAP), would decrease in the closed conformation relative to the open conformation when the peptide was absent, and the bound complex would have less exposed hydrophobics compared to the sum of the exposed hydrophobics of the closed conformation and the peptide alone. We also filtered the bound conformation on interface design metrics, including ddG, cms and SASA.(*33*) This pipeline for designing effector-binding hinges was able to generate very diverse outputs, with large differences in changes in shape and size (Figure S1).

**One-sided two-state design for swapped peptide targets**

To generate swapped-peptide designs, we started from the state X and state Y backbones of cs074, cs201, cs221, and js007, including peptide backbones. Peptide sequences were replaced, using the sequences of cs074B, cs201B, and cs221B. In cases where the peptide backbone was longer than the new peptide sequence, all combinations of N-terminal and C-terminal truncations of the peptide backbone were tested. In cases where the length of the new peptide sequence exceeded that of the backbone, all possible combinations of N- and C-terminal extensions were tested by adding idealized helical residues. All subsequent design steps locked the peptide sequence. The hinge-peptide interface was designed in PyRosetta FastDesign for one repeat with fixed backbone followed by two repeats with flexible backbone, then sequences were improved by design with proteinMPNN using a temperature of 0.2 and model version v_48_020. Structures of proteinMPNN sequences were predicted with AlphaFold2 using the Rosetta design model as initial guess, and designs that predicted with mean pLDDT < 92, RMSD to reference > 1.5, or mean PAE interaction > 5 were discarded. Poses were combined with state X models from the parent hinges, and residues were linked between states for PyRosetta multistate design with flexible backbone followed by MPNN multistate design with the same settings as previous proteinMPNN design. An increased success rate was observed when performing proteinMPNN multistate design with a 60%-40% bias toward state Y sequences. State

Y structures were predicted with AF2-IG, and mean pLDDT, RMSD to reference, and mean PAE interaction cutoffs of 93, 1.5, and 5, respectively. State X structures were also predicted with AlphaFold2 with the same cutoffs, excluding mean PAE interaction. 4 out of 9 possible pairs of parent hinge and peptide sequence produced AlphaFold2-verified models. 20 designs were ordered, expressed, and tested for binding by fluorescence polarization (FP). A common failure mode for these designs was low levels of soluble expression, but 15/19 with sufficient expression for FP displayed detectable binding to the intended peptide. 9 were selected based on soluble expression levels and on-target affinity for further characterization. Only one design (CSW13, Figure S5) was determined to have no off-target binding to the peptides in the parent set, while 4/9 designs still bound the original peptide with higher affinity than the intended one. The remaining 4 bound the intended peptide with the highest affinity, but also displayed binding to one or more other peptides; an example of this is shown in CSW20 (Figure S5)

**Design of 3 helix bundles**

Starting from AF2 models of validated hinge-peptide complexes, we sketched rough 3hb backbones in PyMOL by manually positioning two additional helices to buttress the bound helical effector peptide. For each sketch, we extracted the center four residues of the placed helices and used inpainting with RoseTTAFold to generate 1000 3hb backbones scaffolding those fragments onto the effector peptide. During the inpainting process, residues on the effector peptide interfacing with either of the placed helices were allowed to mutate; this and the placements of the four-residue fragments guided inpainting to build valid 3hb backbones that roughly aligned with the sketches. For the best 10% (by RoseTTAFold pLDDT) of backbones generated from each sketch, sequences were optimized using ProteinMPNN. AF2-IG was used to predict the structure of the designed 3hbs with and without their target hinge, selecting only those designs which retained the same structure in both predictions and bound their target hinge with the same interface as the original effector peptide. We experimentally characterized the 1–3 designs per sketch that showed the best PAE interaction in the bound prediction, pLDDT in both predictions, and structural diversity (by eye).

**Design of hinge-armed trimers**

To fuse hinge cs221 to the asymmetric unit (asu) of a validated C3-symmetric homotrimer(*38*), we manually positioned the two proteins such that they formed a large interface, their termini were near, and the angle of hinge switching was approximately perpendicular to the homotrimer axis of symmetry. We used inpainting with RoseTTAFold to generate 100 loop backbones between the N-terminus of cs221 and the homotrimer asu, allowing residues in the interface between the two proteins to mutate. To improve visibility of the conformational change in nsEM, we extended the C-terminal end of cs221 by fusing it to LHD101B, a previously validated monomeric protein(*39*). Again, we manually positioned the two proteins such that they formed a large interface and their termini were near, then used inpainting with RoseTTAFold to generate 100 loop backbones between those termini, allowing residues in the interface to mutate. For the best 20% (by RoseTTAFold pLDDT) of backbones generated for each fusion, we optimized sequences of the fusion region using ProteinMPNN. We combined the most confidently-predicted (by AF2 pLDDT) LHD101B fusion with each homotrimer asu fusion, modeled each symmetric complex by aligning three copies of each fusion to the original homotrimer, and used AF2-IG to predict the symmetric structure of the designed fusions. We experimentally characterized the 7 most confidently-predicted designs.

**Hinge extension for FRET constructs**

Hinges were extended by aligning a copy of the parent DHR to the first repeat of the hinge and another copy of the parent DHR to the last repeat of the hinge. The extended hinge was then obtained by replacing the first and last repat of the hinge by 2 or more repeats from the parent DHR. For cs221F, the additional repeats were redesigned using proteinMPNN.

**Disulfide stapling**

A custom PyRosetta script was used to identify candidate positions for disulfides that could lock hinges in one conformation. i-j residue pairs where residue i is in domain 1 of the hinge and residue j is in domain 2 of the hinge were exhaustively evaluated using a 6D hashing protocol(*40*). For each candidate pair, 2 separate pdbs were generated for state X and state Y of the hinge with the identified residues i and j mutated to cysteine. AF2-IG was used to filter candidate pairs, selecting only pairs for which the cysteine side chains in the “target” state showed distances and relative orientations compatible with disulfide formations and for which the “off-target” state showed a large distance between cysteine side chains.

**proteinMPNN-based identification of point mutant candidates**

ProteinMPNN was used to generate 100 sequences optimized for state X and another 100 sequences optimized for the state Y-peptide complex. For each state, consensus sequences(*41*) were used to identify non-interface positions with distinct residue preferences that were different between both states. For mutations that AF2 predicted to not affect the global structure, individual protein variants carrying these mutations were experimentally tested using the FP peptide binding assay.

**Cloning, expression, and protein purification**

Genes encoding for proteins and peptides were either purchased as pre-cloned genes from IDT in pet29B expression vectors or purchased as e-blocks from IDT and cloned into custom target vectors using golden gate assembly(*48*). Hinges and 3-helix bundles usually carried a C-terminal SNAC tag(*50*) followed by a 6xHis-tag (Hinge-GSHHWGSTHHHHHH); in some cases the SNAC tag was omitted (Hinge-GSHHHHHH). Peptides were expressed fused to superfolder green fluorescent protein (sfGFP) in either a sfGFP-(linker)-peptide-(linker)-6xHis construct or sfGFP-GSGSENLYFQS-(linker)-peptide-(linker)-6xHis construct. All proteins were expressed either in LEMO21 or NEB BL21(DE3) E. coli cells by autoinduction using TBII media (Mpbio) supplemented with 50x5052, 20 mM $MgSO_4$ and trace metal mix and 50 mg/l Kanamycin. Expression cultures were grown at 37°C for 20-24 h or at 37°C for 5-6 h followed by 24 h at 18°C.

After harvesting with centrifugation, cells were lysed at 4°C with sonication in lysis buffer containing (100 mM Tris HCl pH 8, 200 mM NaCl, 50 mM imidazole, 1 mM PMSF, 1 mM DNase, 1 Pierce™ Protease Inhibitor Mini Tablets, EDTA-free per 100 mL) and clarified with ultracentrifugation at 14-20k x g for 20-40 min. The constructs were bound to ~1 mL Ni-NTA resin (Qiagen) and mixed for 10-60 min. The beads were sequentially washed with 15 mL low salt wash buffer (20 mM Tris HCl pH 8, 200 mM NaCl, 50 mM imidazole), 15 mL high salt wash buffer (20 mM Tris HCl pH 8, 1 M NaCl, 50 mM imidazole), and 15 mL low salt wash buffer. Lysates and buffer were flowed over the resin either using gravity or a vacuum manifold. Proteins were eluted in 1.4 mL of elution buffer (20 mM Tris HCl pH 8, 200 mM NaCl, 500 mM imidazole), after a 0.4 mL pre-elution. In constructs with designed disulfides, copper phenanthroline was then added to the elution at a final concentration of 10 mM, and the resulting mixture was incubated overnight to encourage full formation of the disulfides. In all cases elutions were further purified by SEC/FPLC on Superdex 75 Increase 10/300 GL or Superdex 200 Increase 10/300 GL columns in TBS (20 mM Tris pH 8, 100 mM NaCl), with 0.5 or 1 mL fractionation between 8 and 20 mL. LC-MS was used to confirm correct molecular weight of all purified proteins.

**Protein purification for crystallography**

Constructs were transformed into LEMO21 or NEB BL21(DE3) E. coli and then expressed as 0.5 L cultures in 2L flasks. Proteins were expressed in Studiers M2 autoinduction media with 50 ug/mL kanamycin. Pre-cultures were grown at 37°C for 4 hrs, then 22°C for 14 hr and cultures were inoculated with 10 mL of preculture. Cells were pelleted at 4,000g for 10 minutes, after which the supernatant was discarded. Pellets were resuspended in 40 mL of lysis buffer (100 mM Tris HCl pH 8, 100 mM NaCl, 400 mM imidazole, 1 mM PMSF, 1 mM DNase). Cell suspensions were lysed by microfluidization on a Microfluidics M-100P at 18,000 psi, and the lysate was clarified at 14,000g for 30 minutes. The His-tagged proteins were bound to 8 mL Ni-NTA resin (Qiagen) during gravity flow and washed with 10 mL lysis buffer and 30 mL high salt wash buffer (25 mM Tris HCl pH 8, 1 M NaCl, 40 mM imidazole), then 10mL SNAC cleavage buffer (100 mM CHES, 100 mM Acetone oxime, 100 mM NaCl, 500 mM GnCl, pH 8.6).(*50*) 40 mL SNAC cleavage buffer and 80 uL 1M NiCl2 was added and columns were closed and shook on a nutator for 12 hours in order to cleave. After cleavage the flowthrough was collected and concentrated prior to further purification by SEC/FPLC on a HiLoad 20/600 Superdex 75 pg column in TBS (20 mM Tris pH 8.0, 100 mM NaCl), with 14 mL fractionation between 100 and 290 mL.

**Peptide synthesis**

Peptides were synthesized in-house on a CEM Liberty Blue microwave synthesizer. All amino acids were purchased from P3 Biosystems. Oxyma Pure was purchased from CEM, DIC was purchased from Oakwood Chemical, diisopropyl ethylamine (DIEA) and piperidine were purchased from Sigma-Aldrich. Dimethylformamide (DMF) was purchased from Fisher Scientific and treated with an Aldraamine trapping pack prior to use. 5(6)-carboxytetramethylrhodamine carboxylic acid (5(6)-TAMRA) was purchased from Novabiochem. Synthesis was done on a 0.1 mmol scale on CEM Cl-MPA resin. Five equivalents of each amino acid were activated using 0.1 M Oxyma with 2% (v/v)

DIEA in DMF, 15.4% (v/v) DIC, and coupled twice on resin for 2 min per coupling with microwave irradiation. For TAMRA labeled peptides, peptides were washed with DMF post-synthesis, then incubated for 3h with 5(6)-TAMRA carboxylic acid (3 eq.), HATU (3 eq.), and DIEA (5 eq.) in DMF, then washed with DMF (3x) followed by DCM (3x) to prepare for global deprotection. Global deprotection was accomplished with a TFA/water/TIPS/2,2'-(ethylenedioxy)diethanethiol (92.5:2.5:2.5:2.5) mixture for 3 hours. This deprotection mixture was concentrated *in vacuo* to 2-3mL, then precipitated in 30 mL of ice-cold ethyl ether, centrifuged, and decanted, then washed twice more with fresh ether and dried under nitrogen to yield crude peptide for high pressure liquid chromatography (HPLC) purification. The crude peptide was dried and dissolved in a minimal amount of ACN and water to where the entire crude is soluble. This solution was purified on a Zorbax Stablebond C18 (9.4 x 250mm, 5um) column using an Agilent 1260 Infinity HPLC. A linear gradient of water (0.1% TFA) and increasing ACN (0.1% TFA) was used to purify the crude peptides. UV signal was monitored at 214 nm and all peaks were collected. Peak masses were checked using an Agilent G6230B LC-MS and purity was assessed using a C18 column (Higgins Analytical PROTO 300 C18, 10um, 10 x 250mm) on an analytical Agilent 1260 Infinity II HPLC.

**SEC binding assay**

Individual hinge and sfGFP-fused peptides or 3hb were diluted in 20 mM Tris pH 8, 100 mM NaCl and mixed at approximately 1:1 concentrations. 0.5-1 mL of the resulting samples were injected onto a Superdex 200 Increase 10/300 GL columns and the absorbance at 230 nm was used as a readout for binding. For sfGFP-fused peptides, 473 nm was also used as readout. Mixtures were at a total concentration of 2.5 µM or higher.

**Fluorescence Polarization (FP)**

All FP measurements were performed at 25°C in 96-well plates (Corning 3686) using a Synergy Neo2 plate reader and a 530/590 nm filter cube. The buffer for all FP measurements was 20 mM Tris-HCl, 100 mM NaCl, 0.05 % v/v TWEEN20 at pH 8. Titrations were carried out in 96-well format, with 4 replicates per plate and 24 data points per titration (23 steps of two-fold serial dilution of hinges in the presence of TAMRA-labeled peptide at a constant concentration between 0.1 nM and 1 nM) with a final sample volume of 80 µl per well. Titration plates were incubated overnight at room temperature before measuring to ensure complete equilibration. The polarization signal S (as calculated by the Neo2 software) was fitted to the equation

$$S = S_0 + S_1 * f_{AB}$$

$$f_{AB} = \frac{1}{2B_{tot}}\left(A_{tot} + B_{tot} + K_D - \sqrt{(A_{tot} + B_{tot} + K_D)^2 - 4 * A_{tot} * B_{tot}}\right)$$

where $f_{AB}$ is the fraction of peptide that is bound, $A_{tot}$ is the absolute hinge concentration, $B_{tot}$ is the absolute peptide concentration, $S_0$ is the baseline polarization of free peptide, and $S_1$ is the change in polarization upon complex formation.

For FP kinetics experiments a 2x peptide solution and 8 different 2x hinge solutions at different concentrations were prepared separately. 40 µl of each hinge solution were mixed with 40 µl peptide solution using a multichannel pipet and the measurement was started immediately after mixing. Polarization signals S at each concentration were fitted individually to the equation

$$S = S_0 - S_1 * e^{-k_{app}(t_0+t)}$$

where $S_0$ is the amplitude, $S_1$ is the polarization at equilibrium, $k_{app}$ is the apparent rate constant, $t$ is the time after start of the measurement, and $t_0$ is the dead time between mixing and start of the measurement. For each hinge-peptide pair, the apparent rate constants for 8 different concentrations are fitted to the equation

$$k_{app} = k_{off} + k_{on} * A_{tot}$$

where $k_{off}$ and $k_{on}$ are observed off- and on rates and $A_{tot}$ is the absolute hinge concentration.

**FRET**

AlexaFluor 555 C2 maleimide (donor) and AlexaFluor647 C2 maleimide (acceptor) were purchased from ThermoFisherScientific. Stock solutions at ~5 mM were prepared by dissolving 1 mg of each dye in 200 µl DMSO. Hinge variants containing two cysteines were expressed and purified as described above with the modification that 0.5 mM TCEP was used during lysis, IMAC and SEC, and that the buffer for initial SEC contained 20 mM sodium phosphate (PH 7.0) instead of Tris-HCl. After SEC, 500 µl hinge at a concentration of 50 µM was incubated with 500 µM of a single dy for controls or 250 µM each of two dyes. After 2h incubation at room temperature, samples were purified by SEC using a buffer containing 20 mM Tris-HCl and 100 mM NaCl at pH 8.

The buffer for all FRET measurements was 20 mM Tris-HCl, 100 mM NaCl, 0.05 % v/v TWEEN20 at pH 8. Fluorescence spectra were recorded at room temperature using a FluoroMax spectrometer in a 1 cm x 1 cm cuvette at a sample volume of 3 ml. FRET titrations and kinetics measurements were performed at 25°C in 96-well plates (Corning 3686) using a Synergy Neo2 plate reader. Excitation wavelength was 520 nm and emission wavelength was 665 nm (except for donor-donor controls for which emission wavelength was 555 nm, see Figure S7B).

Titrations were carried out in 96-well format, with 4 replicates per plate and 24 data points per titration (23 steps of two-fold serial dilution of effector (peptide or 3hb) in the presence of double-labeled hinge at a constant concentration of 2 nM) with a final sample volume of 80 µl per well. Titration plates were incubated overnight at room temperature before measuring to ensure complete equilibration. The fluorescence signal was fitted to the equation

$$S = S_0 + sign * S_1 * f_{AB}$$

$$f_{AB} = \frac{1}{2A_{tot}}\left(A_{tot} + B_{tot} + K_D - \sqrt{(A_{tot} + B_{tot} + K_D)^2 - 4 * A_{tot} * B_{tot}}\right)$$

where $f_{AB}$ is the fraction of hinge that is bound, $A_{tot}$ is the absolute hinge concentration, $B_{tot}$ is the absolute peptide concentration, $S_0$ is the baseline fluorescence of free hinge, $S_1$is the change in fluorescence upon complex formation, and $sign =- 1$ for cs 201F (which shows a decrease in FRET upon binding) and $sign = 1$ for cs074F and cs221F (which show an increase in FRET upon binding.

For FRET kinetics experiments a 2x hinge solution and 8 different 2x effector solutions at different concentrations were prepared separately. 40 µl of each effector solution were mixed with 40 µl hinge solution using a multichannel pipet and the measurement was started immediately after mixing. Fluorescence signals S at each concentration were fitted individually to the equation

$$S = S_0 - sign * S_1 * e^{-k_{app}(t_0+t)}$$

where $S_0$ is the amplitude, $S_1$ is the Fluorescence at equilibrium, $k_{app}$ is the apparent rate constant, $t$ is the time after start of the measurement, and $t_0$ is the dead time between mixing and start of the measurement, and $sign =- 1$ for cs 201F (which shows a decrease in FRET upon binding) and $sign = 1$ for cs074F and cs221F (which show an increase in FRET upon binding. For each hinge-peptide pair, the apparent rate constants for 8 different concentrations are fitted to the equation

$$k_{app} = k_{off} + k_{on} * B_{tot}$$

where $k_{off}$ and $k_{on}$ are observed off- and on rates and $B_{tot}$ is the absolute peptide concentration.

**DEER - spin label modeling and site selection**

All spin label modeling and distance distribution predictions were performed using chiLife(*35*) with the off-rotamer sampling method(*51*). For each construct, spin label models were made for every site with at least 50 Å$^2$ solvent accessible surface area (SASA) in both conformational states. Pairwise distance distributions were predicted for all modeled spin labels in both states. Site pairs with the largest earth mover's distance (EMD) between the bound and unbound states were manually inspected and site pairs were selected that were predicted to have minimal interference with peptide binding, and conformational change. Two site pairs were chosen for each construct, one predicted to shift the distance distributions to a larger distance upon interaction with substrate and one predicted to shift to a shorter distance.

**DEER - sample preparation**

Hinge variants carrying two cysteines were purified as described above but with 1 mM TCEP added to the lysis buffer and 0.5 mM TCEP added to an intermediate wash buffer. Directly after elution, 50 µL of 200 mM MTSL solution (in DMSO) was added to the entire 1.3 mL elution. After 1-6 h incubation at room temperature the labeling mixture was sterile filtered and purified by SEC. Successful labeling was confirmed by LC-MS.

Before DEER, 20 µM protein samples were prepared in 20 mM tris, 100 mM NaCl at pH 8.0 in D2O and 20 % d8-glycerol (Cambridge Isotope Laboratories, Inc.) supplemented with 100 µM B-peptide when appropriate. Samples (20 – 40 µL) were transferred to quartz capillaries (Sutter Instruments) with an inner diameter of 1.1 mm and an outer diameter of 1.5 mm, flash frozen with liquid nitrogen and stored at -80 °C.

**DEER - measurements**

All DEER experiments were performed on an ELEXSYS E580 EPR spectrometer (Bruker) at Q-band (~34 GHz) using an EN5107D2 resonator (Bruker). A cryogen free cooling system (ColdEdge) was used to maintain a temperature of 50 K. Shaped pulses were generated using a SpinJet arbitrary waveform generator (Bruker). Observer pulses were 60 ns gaussian pulses with a full width at half maximum (FWHM) of 30 ns performed at approximately the center of the field-swept spectrum. Pump pulsers were 150 ns sech/tanh pulses centered 80 MHz above the observer pulses. Sech/tanh pulses were generated using PulseShape (https://gitlab.com/mtessmer/PulseShape) or EasySpin(*52*) with an excitation bandwidth of 80 MHz and a truncation parameter of 10. All sech/tanh pulses were modified to compensate for resonator performance and transmitter nonlinearity. All experiments used 8-step phase cycling and 8-step $\tau_1$ averaging with 16 ns increments from 400 ns to 528 ns. Pump pulse time steps (Δt) and $\tau_2$ times were chosen on a per sample basis and the values for each sample are reported in Supplementary Table 2. Additional parameters including $t_0$ offsets, shot repetition time, total number of averages, and more are reported in Supplementary Table 2.

DeerLab(*53*) was used to analyze all DEER data to simultaneously fit foreground and background using Tikhonov regularization and compactness regularization(*54*). Akaike information criterion (AIC) and the information complexity criterion (ICC) were used to select regularization parameters for Tikhonov and compactness regularizations respectively. Sample fitting parameters including modulation depth, estimated signal to noise, smoothing and compactness regularization parameters are reported in Supplementary Table 2.

**X-Ray crystallography**

All crystallization experiments were conducted using the sitting drop vapor diffusion method. Crystallization trials were set up in 200 nL drops using the 96-well plate format at 20 ˚C. Crystallization plates were set up using a Mosquito from SPT Labtech, then imaged using UVEX microscopes and UVEX PS-600 from JAN Scientific. Diffraction quality crystals formed for 3hb05 in 0.2 M Lithium sulfate, 0.1 M Na-Phosphate-citrate pH 4.2, 20% PEG 1000; for 3hb12 1.8 M Ammonium citrate tribasic pH 7.0; for cs074AB in 0.2 M Calcium acetate, 0.1 M Na cacodylate pH 6.5, 40% PEG 300; for cs207A in 0.1 M SPG buffer pH 7, 25% (w/v) PEG 1500; for cs207AB in 0.2 M Magnesium sulfate and 20% (w/v) PEG 3350.

Diffraction data was collected at the Advanced Light Source beamlines 8.2.2/8.2.1. X-ray intensities and data reduction were evaluated and integrated using XDS(*55*) and merged/scaled using Pointless/Aimless in the CCP4 program suite(*56*). Structure determination and refinement starting phases were obtained by molecular replacement using Phaser(*57*) using the designed model for the structures. Following molecular replacement, the models were improved using phenix.autobuild(*58*); efforts were made to reduce model bias by setting rebuild-in-place to false, and using simulated annealing and prime-and-switch phasing. Structures were refined in Phenix(*58*). Model building was performed using COOT(*59*). The final model was evaluated using MolProbity(*60*). Data collection and refinement statistics are recorded in Supplementary Table 1. Data deposition, atomic coordinates, and structure factors reported in this paper have been deposited in the Protein Data Bank (PDB), http://www.rcsb.org/ with accession code 8FIH (3hb05), 8FVT (3hb12), 8FIT (cs074AB), 8FIN (cs207A) and 8FIQ (cs207AB).

**Negative stain electron microscopy**

Carbon-coated 400 mesh copper grids (01844-F, TedPella,Inc.) were first glow-discharged using a PELCO easiGlow cleaning System. SEC-purified proteins were diluted to 2 µg/ml with Tris Buffer (100 mM Tris, 40 mM NaCl), and then immediately pipetted onto the glow-discharged grid. The protein solution was allowed to sit on the grid for 30s, before being blotted away with Whatman filter paper. 3 uL of 2% uranyl formate stain was added to the grid and then blotted away after 10s. A second and third wash of UF stain was added to the grid, allowed to sit for 10s and 30s respectively, before being blotted away. The grid was allowed to air-dry for 5 minutes. Dried grids were then imaged using a FEI Talos L120C TEM (FEI Thermo Scientific, Hillsboro, OR) equipped with a 4K × 4K Gatan OneView camera, at a magnification of 57,000x and pixel size of 2.49 Å. Once a grid-square with satisfactory stain thickness and contrast was identified, EPU software was used to automatically collect 200-400 micrographs across the square. Micrographs were imported into and analyzed using cryoSPARC v4.0.3. 50-100 particles were manually picked and subjected to 2D classification to find coarse 2D averages that could be used as templates for automated picking of thousands of particles across all micrographs. After automated picking and particle extraction from micrographs, a further round of 2D classification was done to find higher resolution averages of the hinge-bearing cyclic ring proteins in various states and orientations.

#### Supplementary Note 1: Kinetic model for peptide-binding hinges

Our kinetic model comprises the three states X, Y, and YP (state Y bound to the peptide), assuming that a state X bound to the peptide would be sterically unfeasible. The model considers four microscopic rate constants: $k_1$ and $k_{-1}$ describe the conformational change from X to Y and from Y to X, respectively; $k_2$ and $k_{-2}$ respectively describe the association and dissociation of state Y and the peptide. We assume that direct transitions from state X to the complex YP or from YP to X do not occur, but always involve Y as intermediate. Given the model of two coupled equilibria as shown in Figure 5B, the fundamental rate laws for the individual states are

$$\frac{d[X]}{dt} = k_{-1}[Y] - k_1[X] \quad (1)$$

$$\frac{d[Y]}{dt} = k_1[X] - k_{-1}[Y] - k_2[Y][P] + k_{-2}[YP] \quad (2)$$

$$\frac{d[YP]}{dt} = k_2[Y][P] - k_{-2}[YP] \quad (3)$$

With [X], [Y], [P] and [YP] being the concentrations for hinge in state X, hinge in state Y, free peptide, and state Y - peptide complex, respectively.

In our FRET system the measured intensity I (acceptor emission upon donor excitation) can be described as

$$I = [X]I_X + ([Y] + [YP])I_Y = [X]I_X + ([H]_{total} - [X])I_Y$$

$$I = [H]_{total}I_Y + [X](I_X - I_Y) \quad (4)$$

With $[H]_{total}$ = [X] + [Y] + [YP] and assuming one intensity $I_X$ for hinges in state X and another intensity $I_Y$ for hinges in state Y or the state Y-peptide complex.
We performed FRET kinetics measurements using a constant concentration of labeled hinge $[H]_{total}$ and varying peptide concentrations $[P]_{total}$ = [P] + [YP] that were at least 10-fold higher than the hinge concentration ($[P]_{total} >> [H]_{total}$). The resulting data can be fit with a single exponential equation

$$I(t) = c_1 + c_2 e^{-k_{app,FRET}\, t}$$

With constants $c_1$ and $c_2$ and an apparent rate constant $k_{app,FRET}$. Substituting I from (4) gives

$$[H]_{total} I_Y + [X](I_X - I_Y) = c_1 + c_2 e^{-k_{app,FRET} t}$$

Elimination of constant terms leaves

$$[X](I_X - I_Y) = c_2 e^{-k_{app,FRET} t}$$

$$[X] = \frac{c_2}{(I_X - I_Y)} e^{-k_{app} t}$$

Which leads to

$$\frac{d[X]}{dt} = -k_{app,FRET} \frac{c_2}{(I_X - I_Y)} e^{-k_{app,FRET} t} = -k_{app,FRET} [X]$$

Using the fundamental rate law (1) we get

$$-k_{app,FRET} [X] = k_{-1} [Y] - k_1 [X]$$

Solving for $k_{app,FRET}$ gives

$$k_{app,FRET} = k_1 - \frac{[Y]}{[X]} k_{-1}$$

Which shows that the ratio [Y]/[X] is constant over time, meaning that X and Y are in a pre-equilibrium with exchange rates that are faster than the association rates we observe.

In our FP system, we measure polarization values (pol) that depend on the ratio of bound peptide:

$$pol = \frac{[P]}{[P]_{total}} pol_{free} + \frac{[YP]}{[P]_{total}} pol_{bound} = \frac{[P]_{total} - [YP]}{[P]_{total}} pol_{free} + \frac{[YP]}{[P]_{total}} pol_{bound}$$

$$pol = pol_{free} + (pol_{bound} - pol_{free}) \frac{[YP]}{[P]_{total}}$$

with polarization values $pol_{bound}$ and $pol_{free}$ for bound and unbound peptide, respectively and with $[P]_{total}$ = [P] + [YP]. The observed change in polarization upon binding can thus be described as

$$\frac{d[pol]}{dt} = \frac{d[YP]}{dt}\,\frac{pol_{bound} - pol_{free}}{[P]_{total}}$$

Using $[P]_{total}$ = [P] + [YP] the fundamental rate law (3) can be rewritten as

$$\frac{d[YP]}{dt} = k_2\,[Y][P] - k_{-2}\,[YP] = k_2\,[Y]\,([P]_{total} - [YP]) - k_{-2}\,[YP]$$

$$\frac{d[YP]}{dt} = k_2[Y][P]_{total} - (k_2[Y] + k_{-2})\,[YP] \qquad (5)$$

Given a large excess of hinge over peptide we can assume that

$$[X] + [Y] = [H]_{total} - [YP] \approx [H]_{total}$$

$$[Y] = F_Y\,[H]_{total} \qquad (6)$$

with $F_Y$ defined as the fraction of hinge that is in state Y in equilibrium:

$$F_Y = \frac{[Y]}{[X]+[Y]} = \frac{K_{XY}}{K_{XY}+1}$$

In equilibrium, $[YP] = [YP]_{eq}$ and $\frac{d[YP]}{dt} = 0$ which gives

$$\frac{d[YP]}{dt}(eq) = k_2[Y][P]_{total} - (k_2[Y] + k_{-2})\,[YP]_{eq} = 0$$

$$k_2[Y][P]_{total} = (k_2[Y] + k_{-2})\,[YP]_{eq} \qquad (7)$$

Substituting (7) in (5) gives

$$\frac{d[YP]}{dt} = k_2[Y][P]_{total} - (k_2[Y] + k_{-2})\,[YP] = (k_2[Y] + k_{-2})\,[YP]_{eq} - (k_2[Y] + k_{-2})\,[YP]$$

$$\frac{d[YP]}{dt} = (k_2[Y] + k_{-2})\,(\,[YP]_{eq} - [YP]) \qquad (8)$$

The reaction approaches equilibrium with

$$\frac{d([YP]_{eq} - [YP])}{dt} = -\frac{d[YP]}{d[t]} = -(k_2[Y] + k_{-2})([YP]_{eq} - [YP]) \quad (9)$$

Defining the apparent rate constant for association using (6) as

$$k_{app} = k_2[Y] + k_{-2} = k_2F_Y[H]_{total} + k_{-2} \quad (10)$$

we can rewrite (9) as

$$\frac{d([YP]_{eq} - [YP])}{dt} = k_{app}([YP]_{eq} - [YP])$$

Solving the differential equation gives the displacement from equilibrium $[YP]_{eq}$ - [YP]:

$$([YP]_{eq} - [YP])(t) = c\,e^{k_{app}t}$$

$$[YP](t) = [YP]_{eq} - c\,e^{k_{app}t}$$

Substituting $[YP](0) = [YP]_0$ at $t = 0$ gives $c = [YP]_{eq} - [YP]_0$ and subsequently

$$[YP](t) = [YP]_{eq} - ([YP]_{eq} - [YP]_0)\,e^{k_{app}t}$$

If $[YP]_0 = 0$ as in our kinetic experiments we get

$$[YP](t) = [YP]_{eq} - [YP]_{eq}\,e^{k_{app}t} \quad (11)$$

Single-exponential fits of our FP kinetics experiments show $k_{app}$ to increase linearly with the total hinge concentration, which we can fit using (10):

$$k_{app} = k_2F_Y[H]_{total} + k_{-2} = k_{on}[H]_{total} + k_{-2}$$

With an observed on rate $k_{on} = k_2F_Y$

If we assume that variants of a given hinge, such as mutants or stapled versions, have the same microscopic on rate, $k_{2,}$ we can estimate $F_y$ by comparing observed on rates for variants a and b:

$$\frac{F_{Y,a}}{F_{Y,b}} = \frac{k_{on,a}}{k_{on,b}}$$

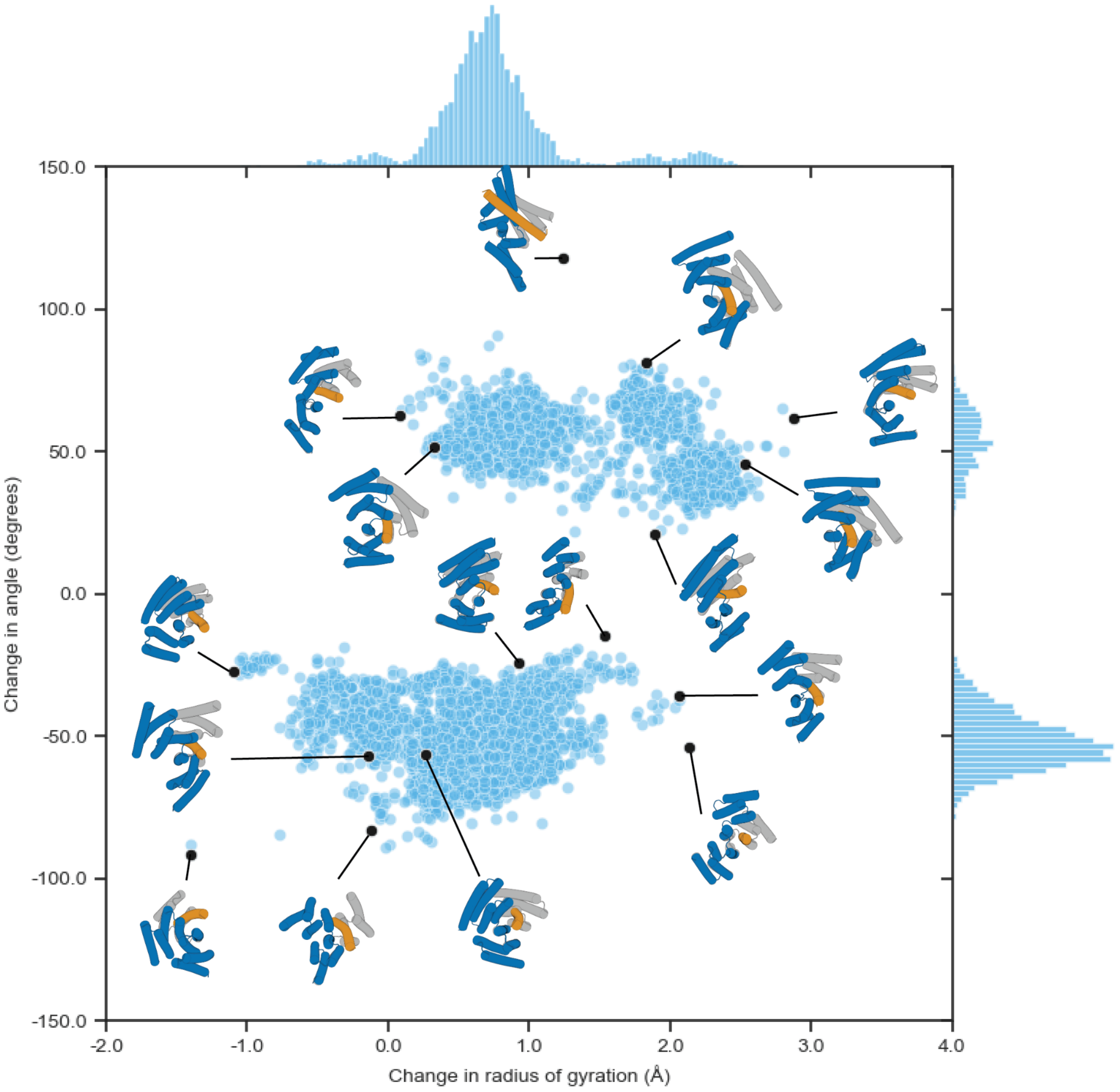


**Figure S1: Diversity of hinge structures and conformational changes.** Light blue points represent individual designs, with positions as a function the angle change of the hinge (state Y - state X, N-terminus-midpoint-C-terminus angle), measured in degrees, and the change in radius of gyration of the hinge (state Y - state X) in angstrom as computed by PyRosetta. Black points are representative examples and are depicted as cartoon models (state X in gray, state Y in blue and orange).

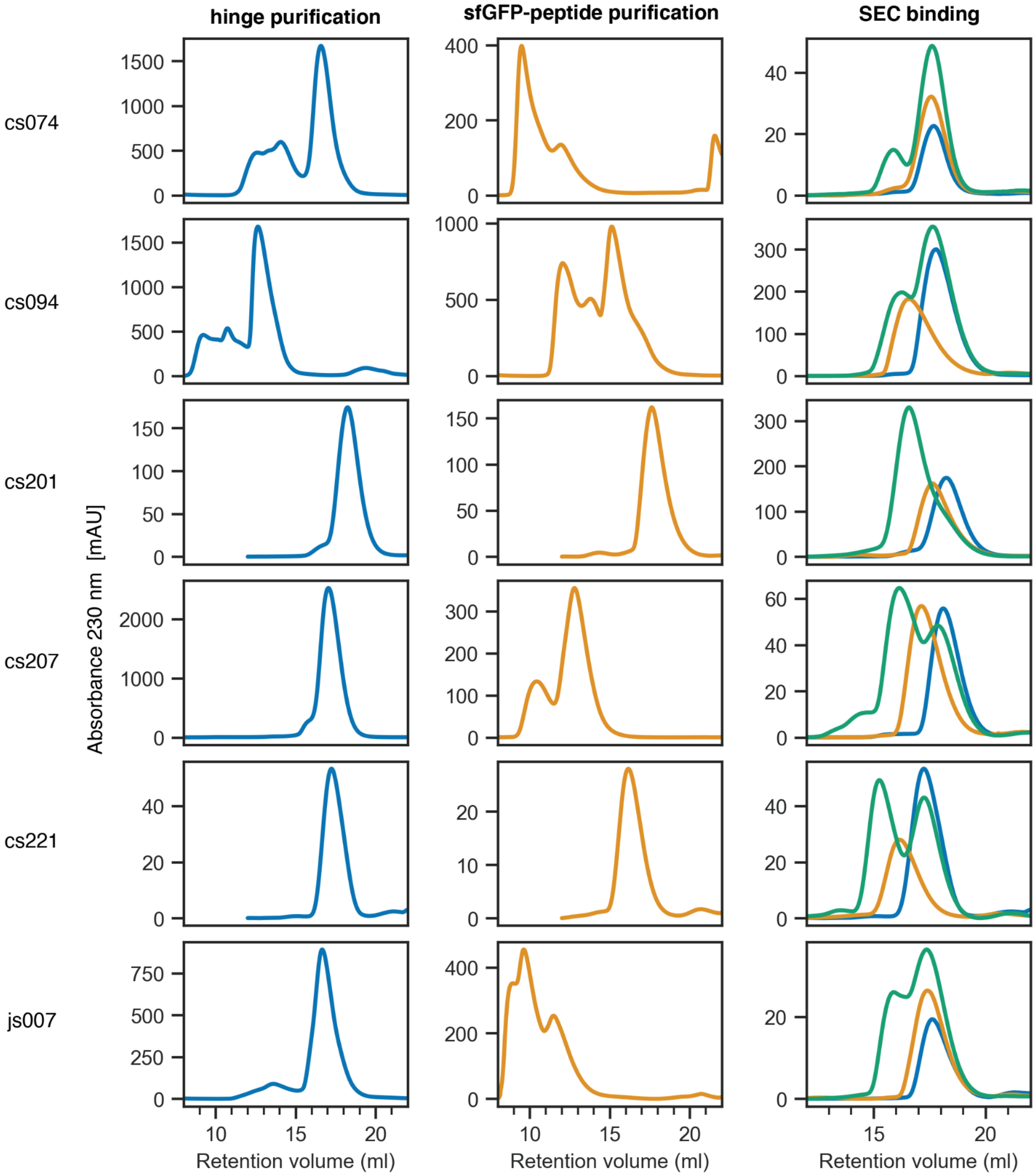


**Figure S2: Size exclusion chromatography (SEC) of the hinges shown in Figure 2.** Purification runs of hinges (left, blue) and sfGFP-peptide fusions (center, orange) were performed on Superdex 75 Increase 10/300 GL columns (Cytiva). SEC binding experiments (blue: hinge, orange: sfGFP-peptide fusion, green: mixture of both) were performed on Superdex 200 Increase 10/300 GL columns (Cytiva).

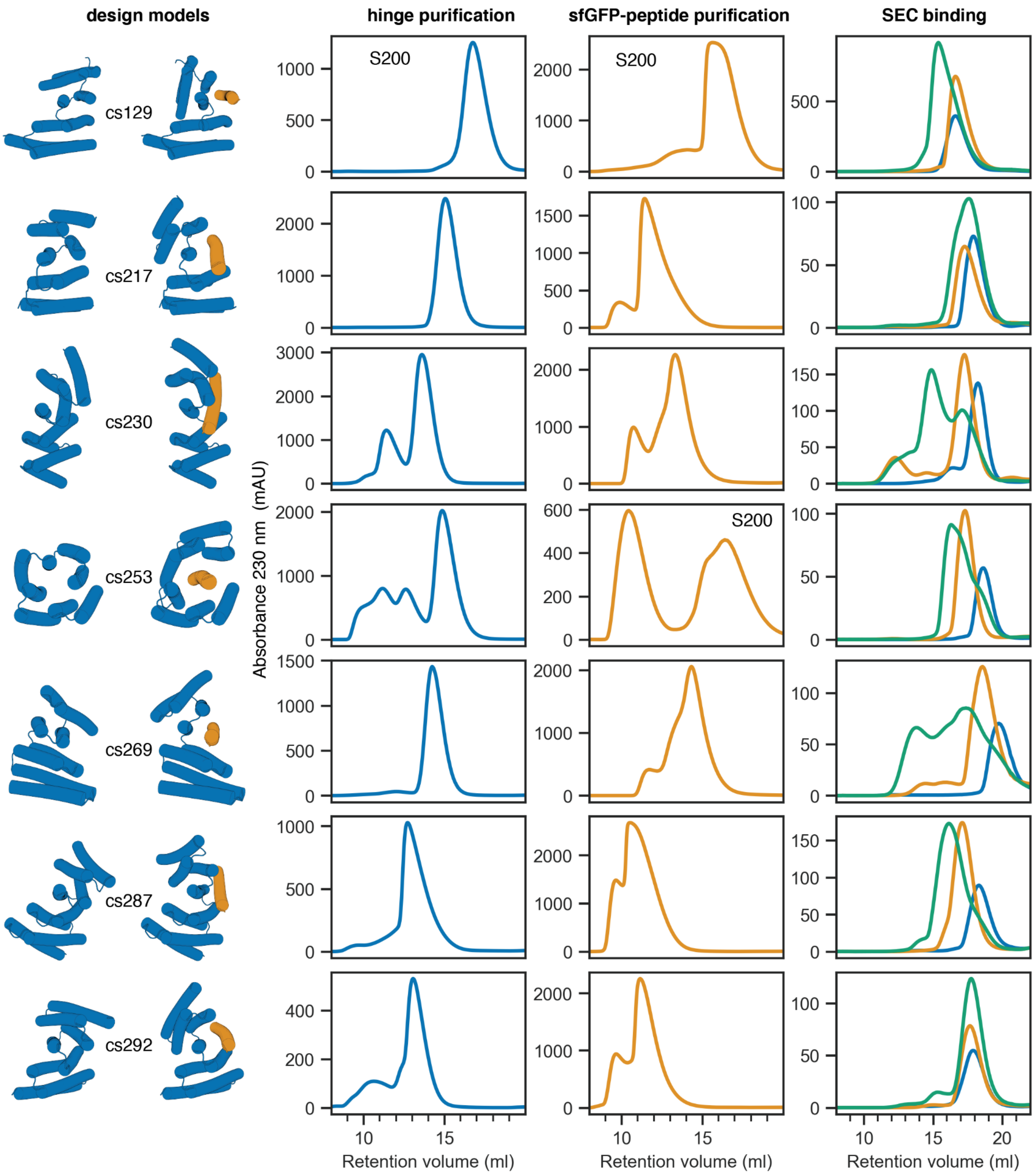


**Figure S3: SEC Characterization of additional hinges not shown in Figure 2.** Purification runs of hinges (left, blue) and sfGFP-peptide fusions (center, orange) were performed on Superdex 75 Increase 10/300 GL columns (Cytiva) except for traces with label “S200” that were run on a Superdex 200 Increase 10/300 GL column. SEC binding experiments (blue: hinge, orange: sfGFP-peptide fusion, green: mixture of both) were performed on Superdex 200 Increase 10/300 GL columns (Cytiva).

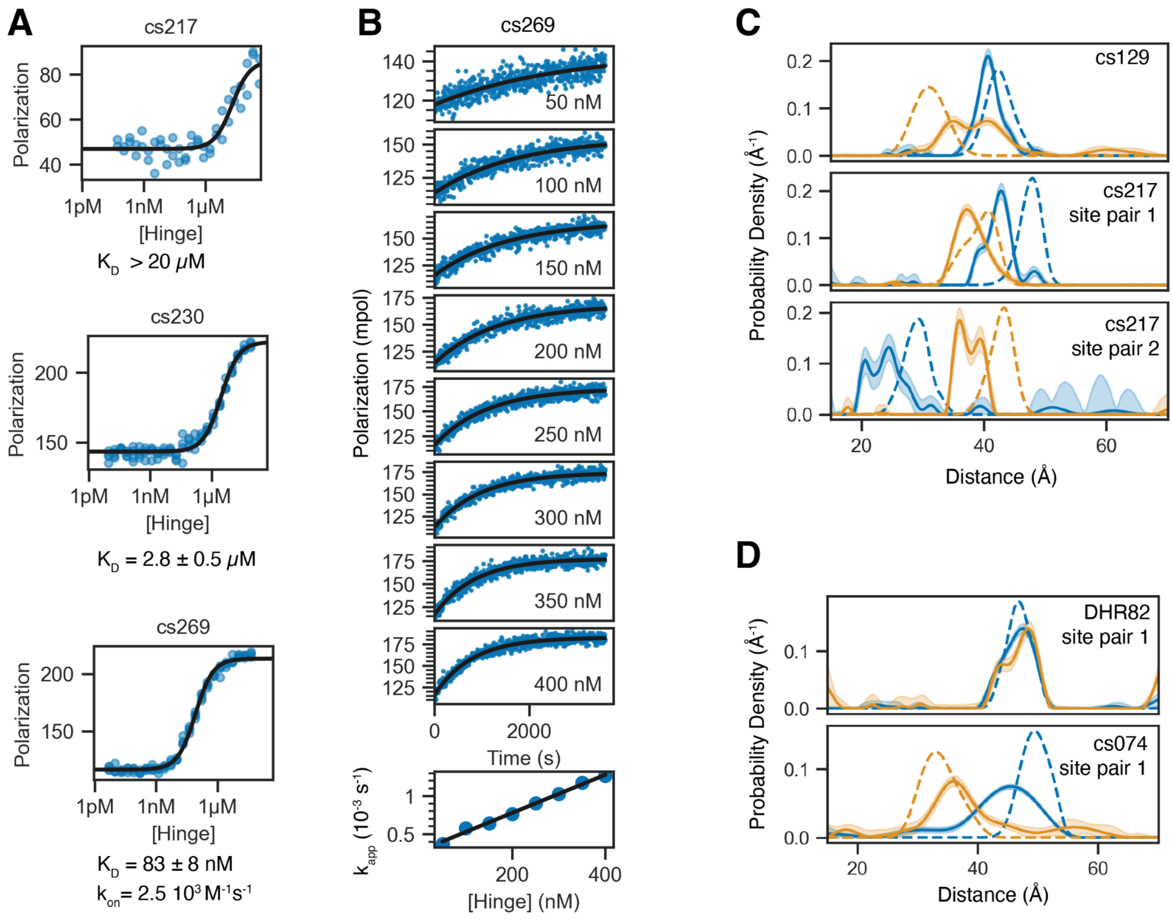


**Figure S4: Additional characterization experiments. A)** FP-based titration using TAMRA-labeled peptide at 1 nM and varying concentrations of hinge. For cs217 the signal never reached a plateau, thus only a lower bound for the $K_D$ can be estimated. **B)** FP-based kinetics experiment using 5 nM TAMRA-labeled peptide cs269B and varying concentrations of hinge cs269 as indicated by plot labels. Top: All kinetic traces were fitted using a single-exponential equation (black lines). Bottom: Apparent rate constants (blue points) from the single exponential fits plotted against the hinge concentration and fitted as linear (black lines). The slope of the linear fit gives the observed on rate $k_{on}$. **C)** DEER distance distributions for hinges cs129 (one site pair) and cs217 (two site pairs) in absence of peptide (blue) and with excess peptide (orange). Dashed lines are simulated distributions based on design models, solid lines are fits to the experimental data, shaded areas are confidence intervals of these fits. **D)** DEER control experiment using DHR82 (the parent of hinge cs074) labeled at the same sites as cs074. Top: DHR82 shows a sharp peak that does not shift upon addition of the peptide cs074B. Bottom: The distance distribution for cs074 that is shown in Figure 2 is shown again as comparison to the DHR82 distribution. The hinge cs074 in absence of peptide shows a slightly broader peak than the parent DHR, suggesting increased flexibility and conformational breathing.

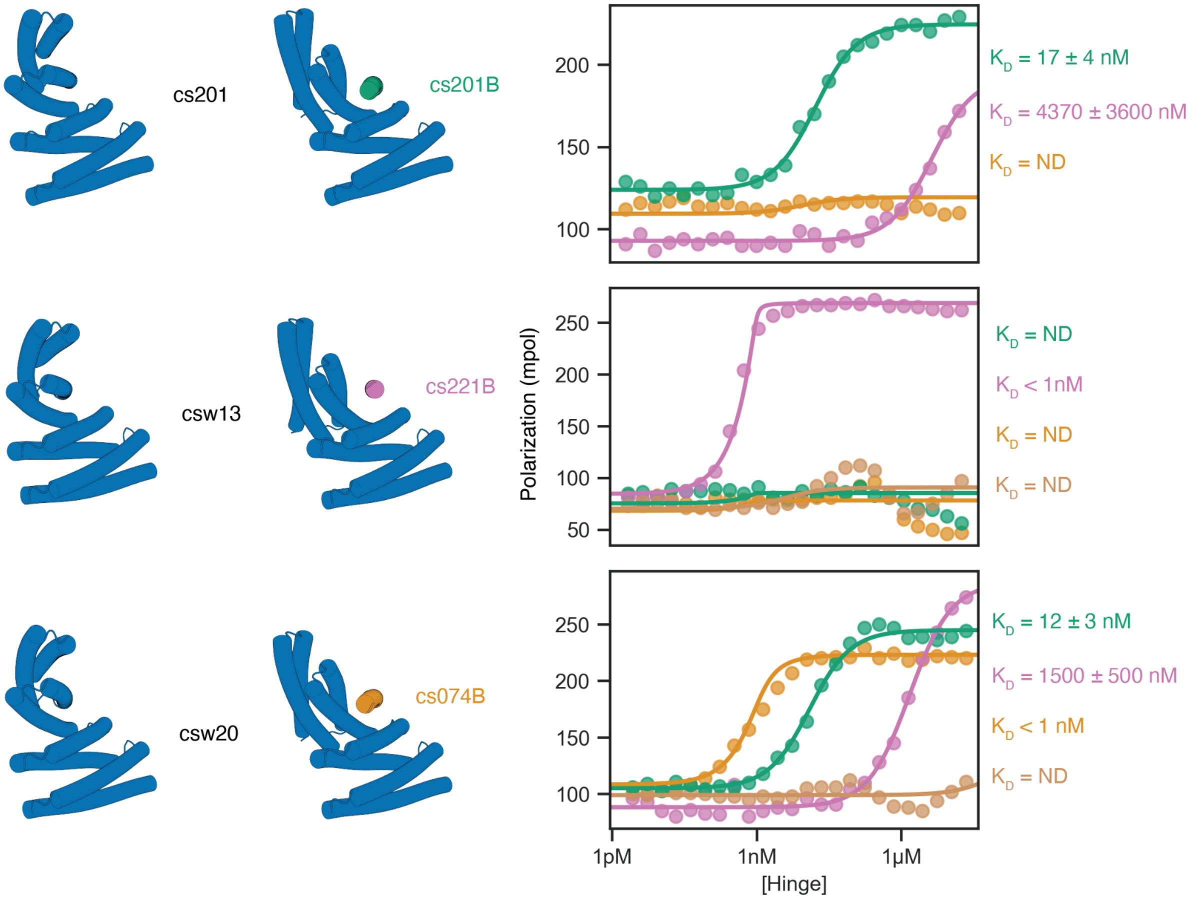


**Figure S5: Redesign of hinges to bind other target peptides.** An experimentally tested hinge (top, cs201) is redesigned using a one-sided two-state design approach to bind to different target peptides. Hinges csw13 and csw20 are designed to bind to peptides cs221B and cs074B, respectively, while having a backbone conformation that is similar to the parent cs201 that binds cs201B. Left: design models, right: FP titrations using 1 nM peptide (green: cs201B, pink: cs221B, orange: cs074B, brown: js007B) and varying concentrations of hinge (from top to bottom: cs201, csw13, csw20). csw13 is an example of a successful orthogonal redesign that specifically binds the new target peptide cs221B while not binding the parent target peptide cs201B or the off-target peptides cs074B or js007B. csw20 is a less orthogonal example that binds the new target peptide cs074B most strongly but still binds to the parent-target peptide cs201B and to the off-target peptide cs221B albeit weaker than the new target.

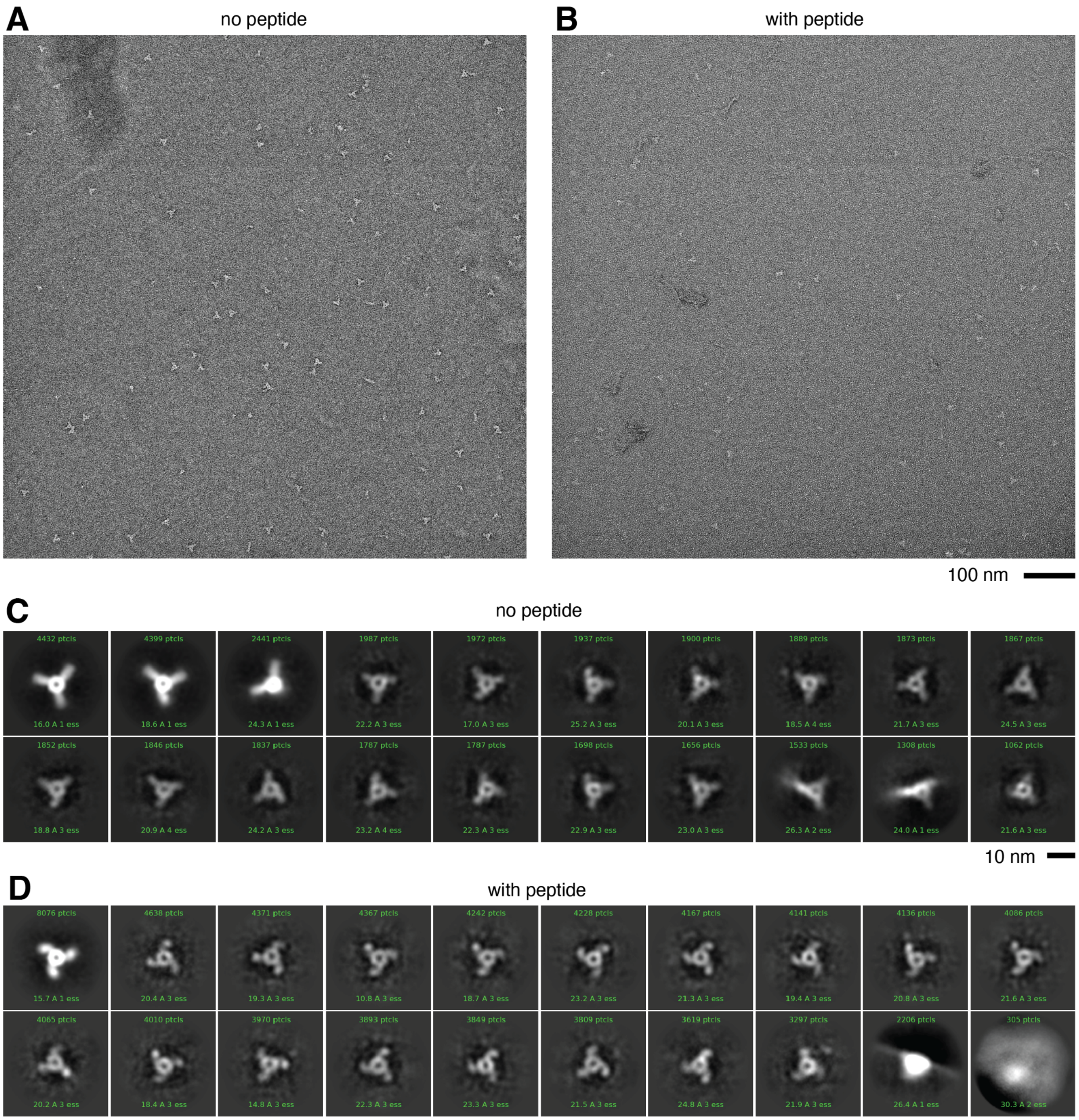


**Figure S6: negative stain electron microscopy on hinge-armed trimers.**
**A,B)** Field-of-view electron micrographs of hinge-armed trimers in absence (A) or presence (B) of peptide cs221B. **C,D)** Class averages from one round of classification (20 classes) using particles obtained from hinge-armed trimers in absence (C) or presence (D) of peptide cs221B.

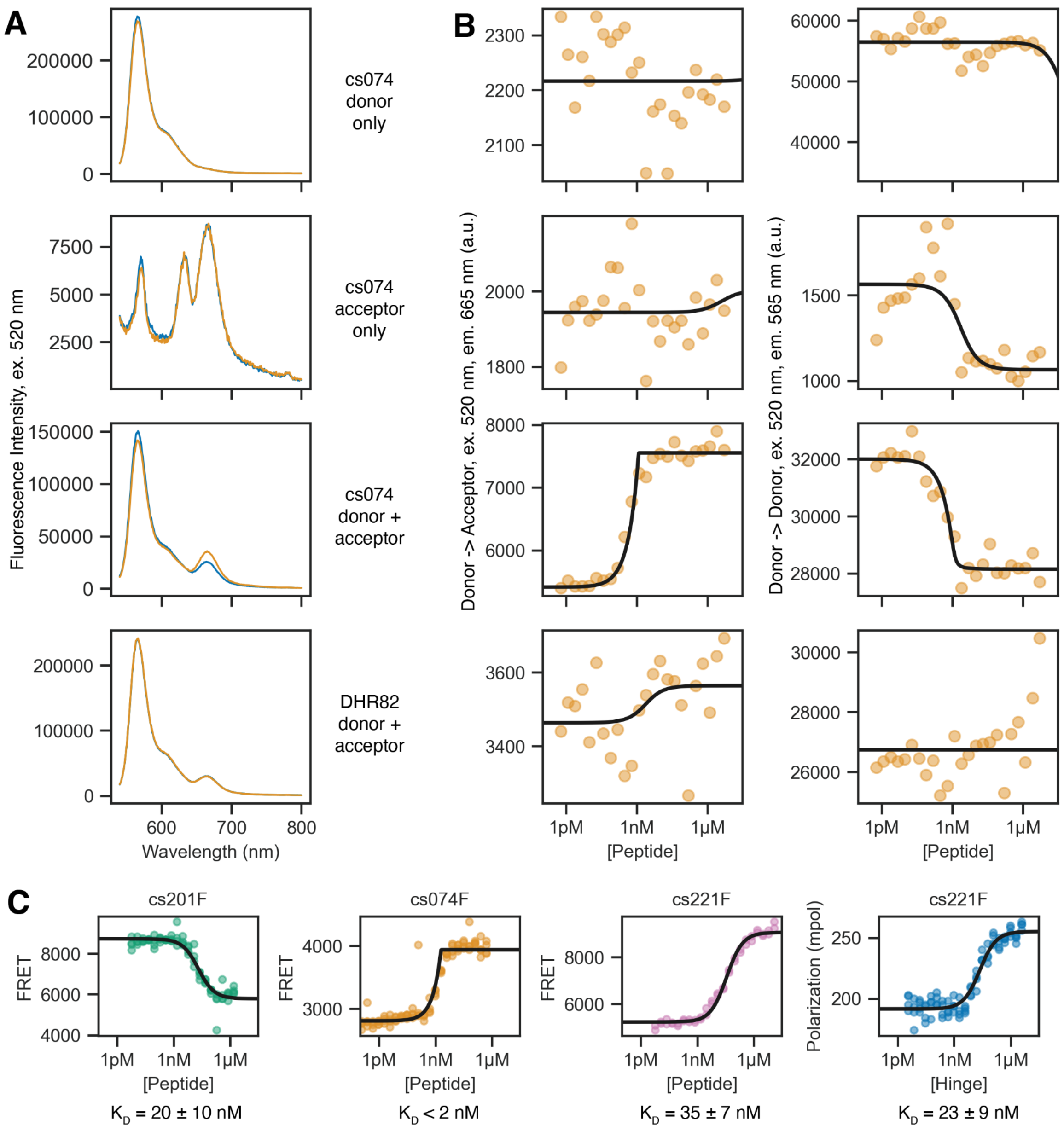


**Figure S7: Additional FRET experiments. A)** Fluorescence spectra of proteins (2nM) without (blue) or with (orange) peptide cs074B (5 nM for cs074 variants, 2 μM for DHR82). Proteins from top to bottom: cs074F labeled with AlexaFluor 555 (= donor only), cs074F labeled with AlexaFluor 647 (= acceptor only), cs074F labeled with a 1:1 mixture of AlexaFluor 555 and AlexaFluor 647 (= donor+acceptor), DHR82 labeled with a 1:1 mixture of both dyes. DHR82 shows no significant change in FRET upon addition of the peptide. **B)** Titrations of the same proteins as in A at 1.2 nM and peptide cs074B at varying concentrations. Left: Acceptor emission upon donor excitation, right: donor emission upon donor excitation. **C)** Replicate titrations of the FRET-labeled extended hinges shown in Figure 4 (2 nM hinge), and FP titration of the unlabeled extended hinge cs221F (right, 1 nM TAMRA-peptide cs221B).

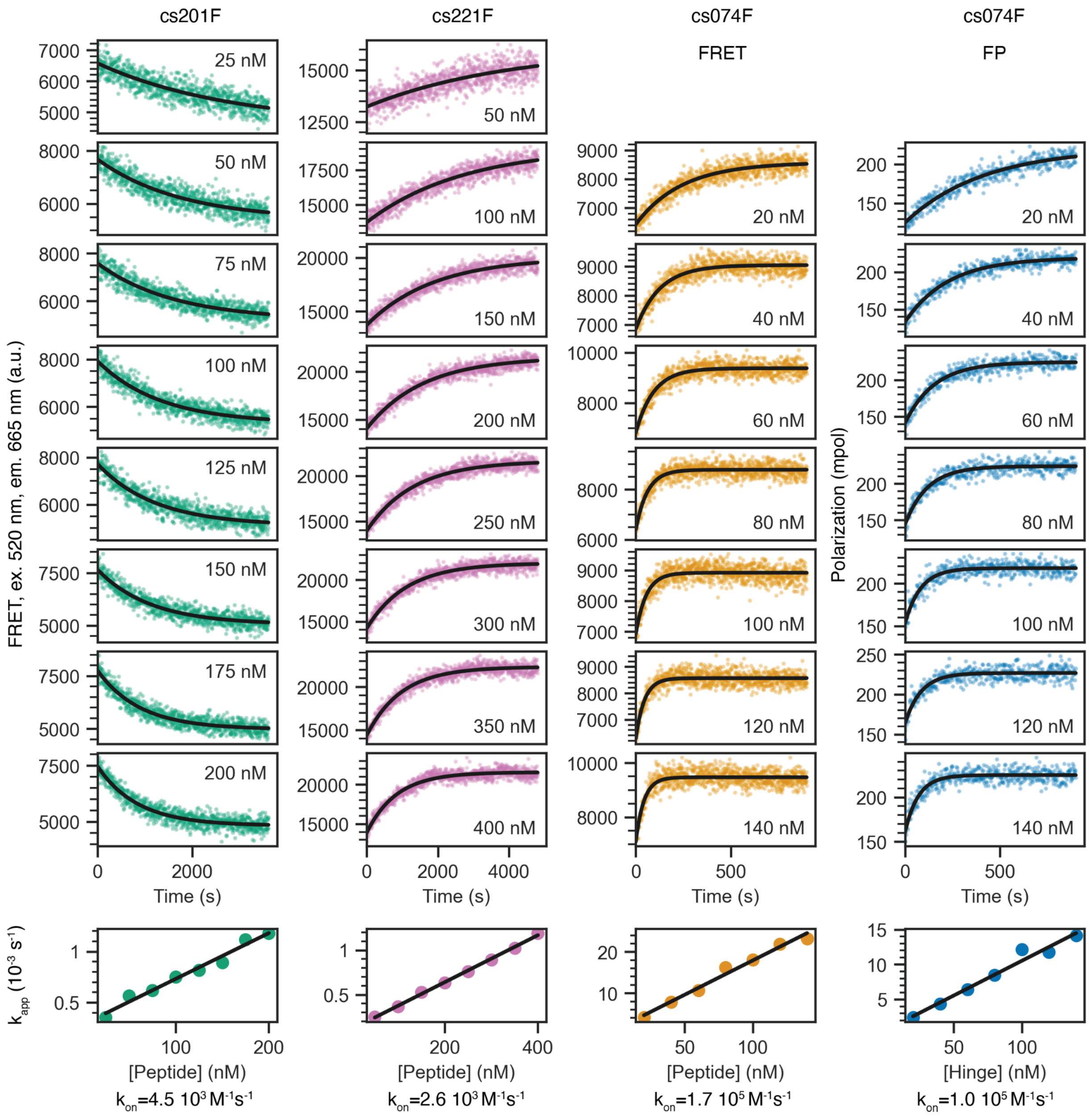


**Figure S8: Full kinetics measurements of the extended hinges shown in Figure 4.** Columns 1-3: FRET kinetics using extended hinges labeled with AlexaFluor 555 and AlexaFluor 647 at a constant concentration (2 nM for cs201F and cs074F, 5 nM for cs221F) and corresponding peptides at varying concentrations. Column 4: FP kinetics using TAMRA-labeled peptide cs074B at 2nM and extended hinge cs074F at varying concentrations. All kinetic traces (rows 1-8) were fitted using a single-exponential equation (black lines). Row 9 shows apparent rate constants from the single exponential fits plotted against the hinge concentration and fitted as linear (black lines). The slope of the linear fit gives the observed on rate $k_{on}$.

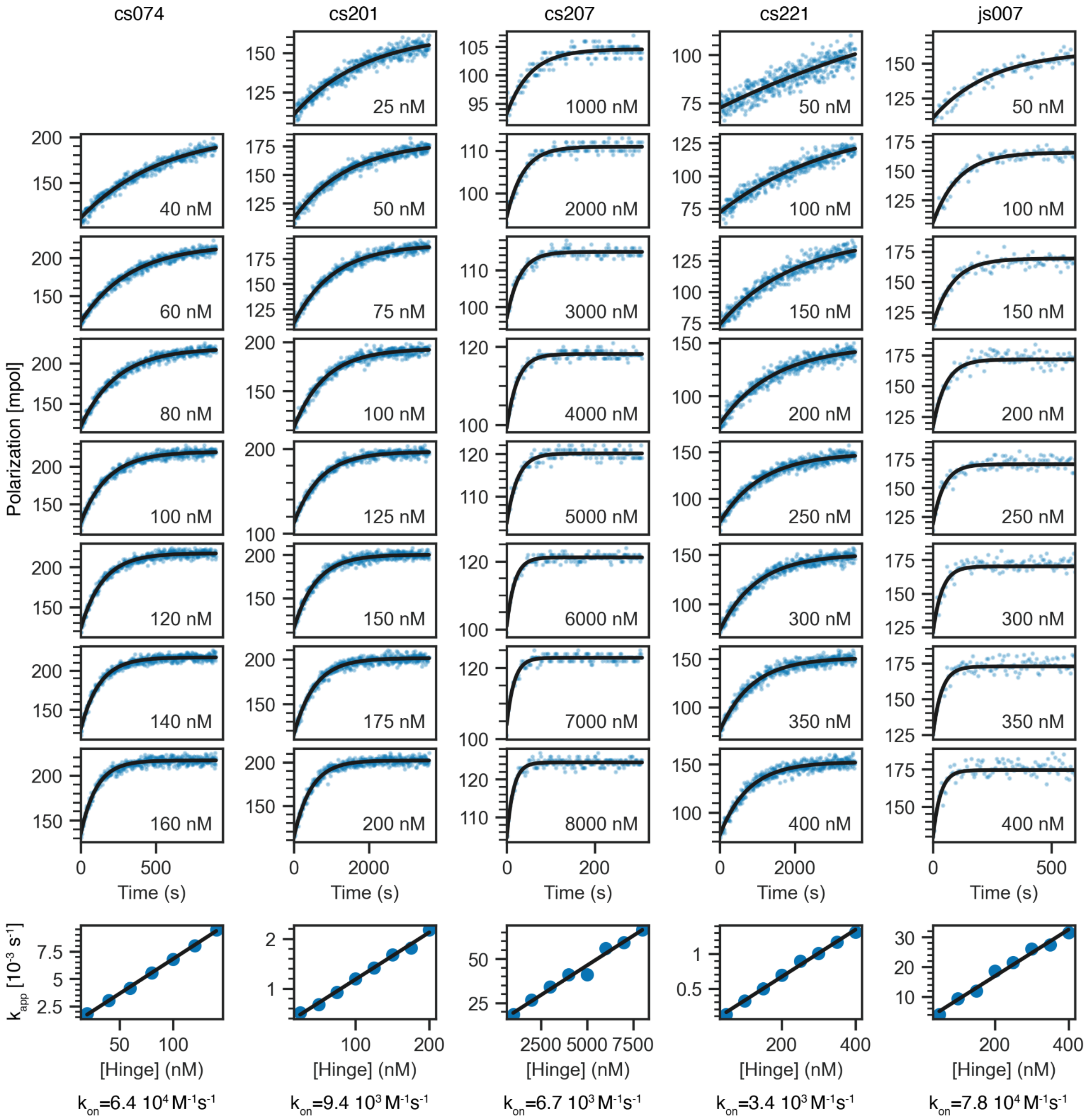


**Figure S9: FP kinetics measurements of hinges shown in Figure 2.** Rows 1-8: TAMRA-labeled peptide at a constant concentration (2 nM for cs074, 50 nM for cs207, 5 nM for cs201, cs221, and js007) was mixed with hinge at varying concentrations (labels in each plot indicate the hinge concentration for the corresponding experiment). All kinetic traces were fitted using a single-exponential equation (black lines). Row 9: Apparent rate constants from the single exponential fits plotted against the hinge concentration and fitted as linear (black lines). The slope of the linear fit gives the observed on rate $k_{on}$.

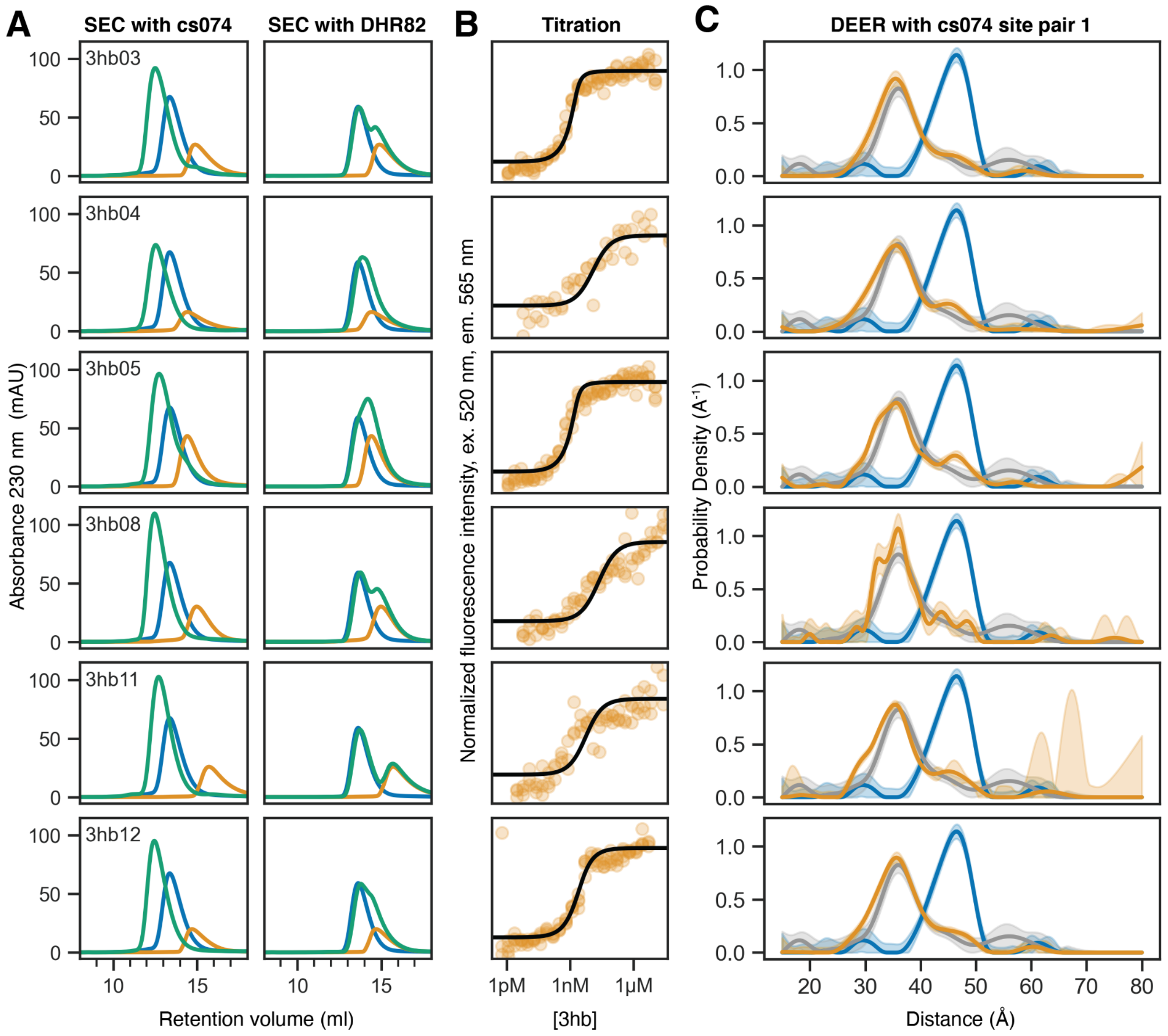


**Figure S10: Additional three-helix bundles that bind cs074. A)** SEC binding experiments. Left: Overlaid chromatograms of three-helix bundles (orange), hinge cs074 (blue) and mixtures of both (green) show clear binding. Right: Overlaid chromatograms of three-helix bundles (orange), parent DHR82 (blue) and mixtures of both (green) show no significant binding. **B)** FRET-based titration experiments using 2 nM labeled cs074F and varying concentrations of 3hb show that the 3hb designs bind to the target hinge with nanomolar affinities and cause a conformational change. **C)** DEER experiments with MTSL-labeled cs074 show that the 3hb designs cause the same conformational change as the original peptide cs074B. Blue: hinge only, gray: hinge + peptide cs074B, orange: hinge + 3hb. Shaded areas indicate 95% confidence intervals.

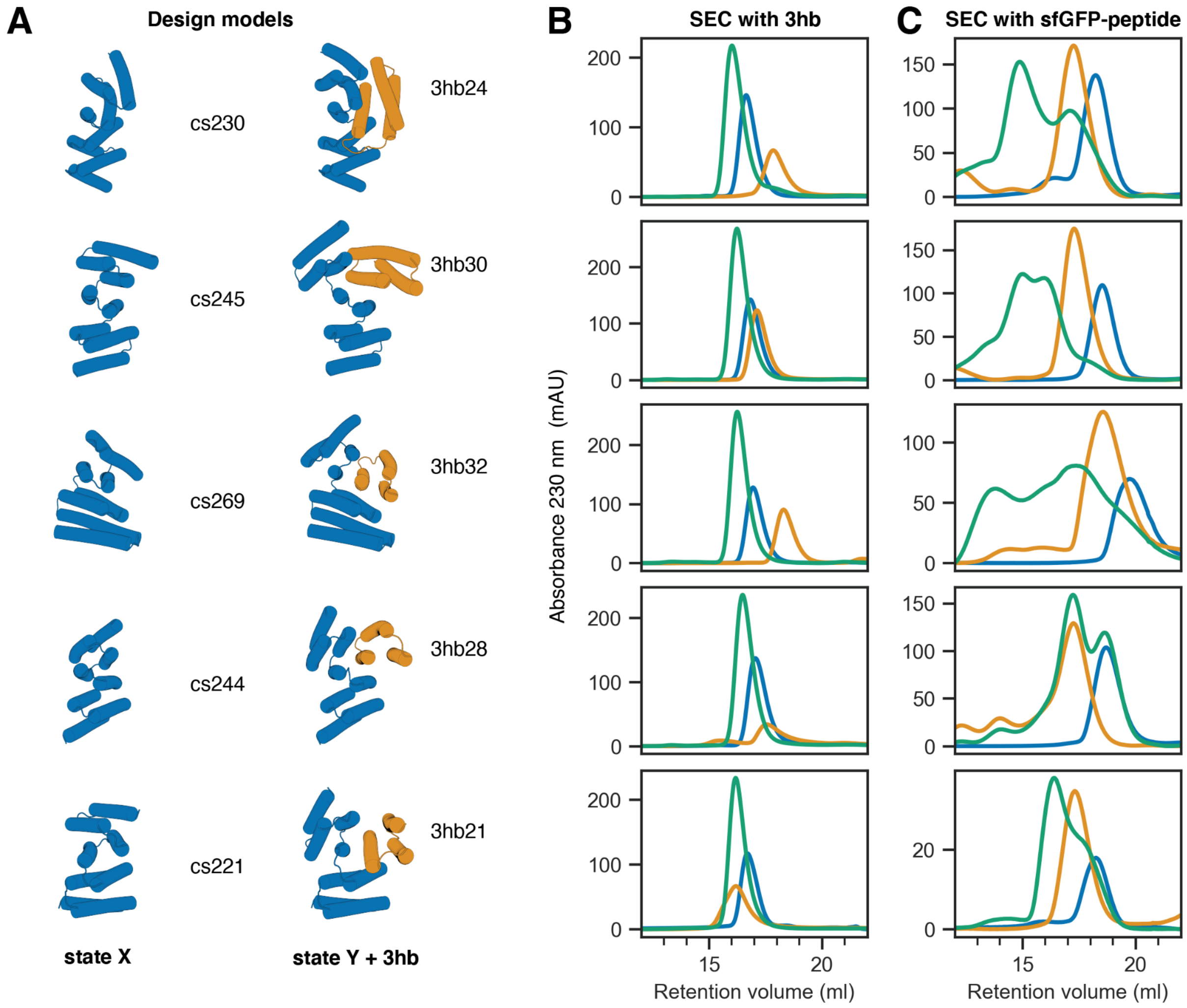


**Figure S11: Additional three-helix bundles. A)** Models of hinges (blue) in state X and in state Y bound to a three-helix bundle (3hb, orange). **B)** SEC binding experiments of hinge (blue), 3hb (orange), and mixture of both (green) show clear monodisperse complex peaks. **C)** SEC binding experiments of the same hinges with the original peptides fused to superfolder green fluorescent protein (sfGFP). For cs230, cs245, and cs269 the hinge-peptide complex shows higher-order peaks while the corresponding hinge-3hb peaks look much cleaner. For cs244, the original peptide showed no clear binding in the SEC experiment, while 3hb28 shows clear binding.

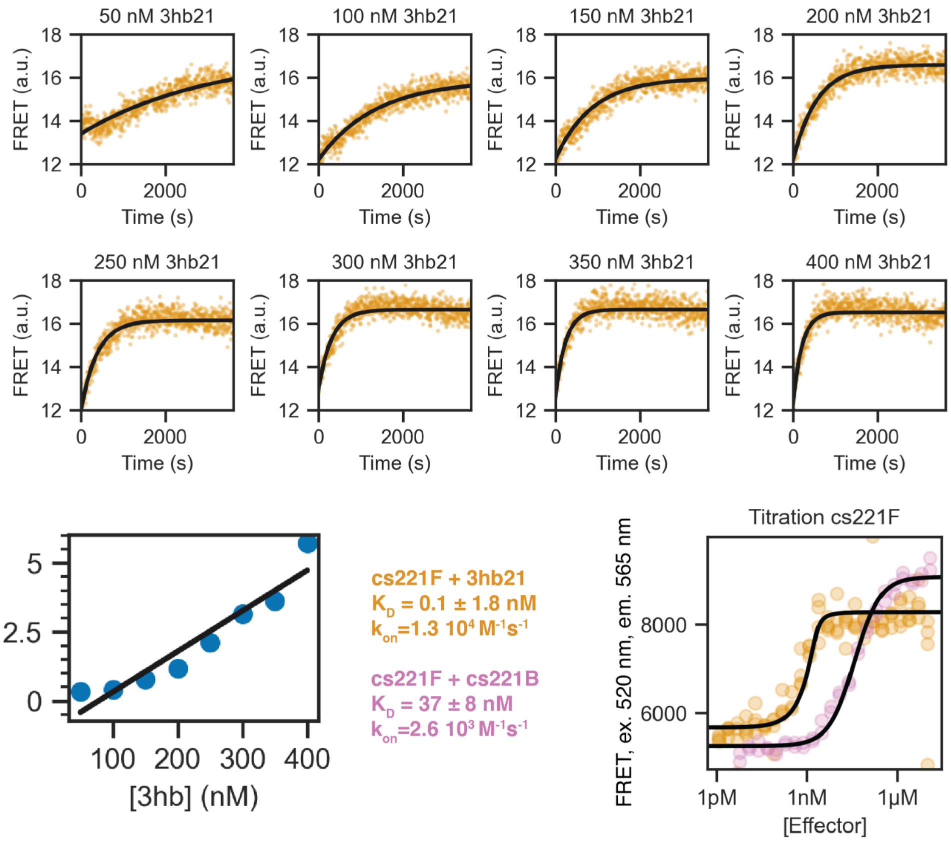


**Figure S12: FRET-based quantitative analysis of the interaction between cs221F and 3hb21.** Individual kinetic traces were obtained using a constant hinge concentration of 5 nM and varying 3hb concentrations as indicated by plot labels. Single exponential fits give apparent rate constants that increase linearly with the total 3hb concentration (orange points in pseudo-first order plot).The linear fit of $k_{app}$ against 3hb concentration gives an observed on rate that is 5 times faster than the observed on rate of the original peptide (pink points). FRET titration of 2 nM hinge cs221F and varying concentrations of 3hb (orange) gives a KD below 2 nM which is at least 20 times stronger than the KD of the original peptide(pink).

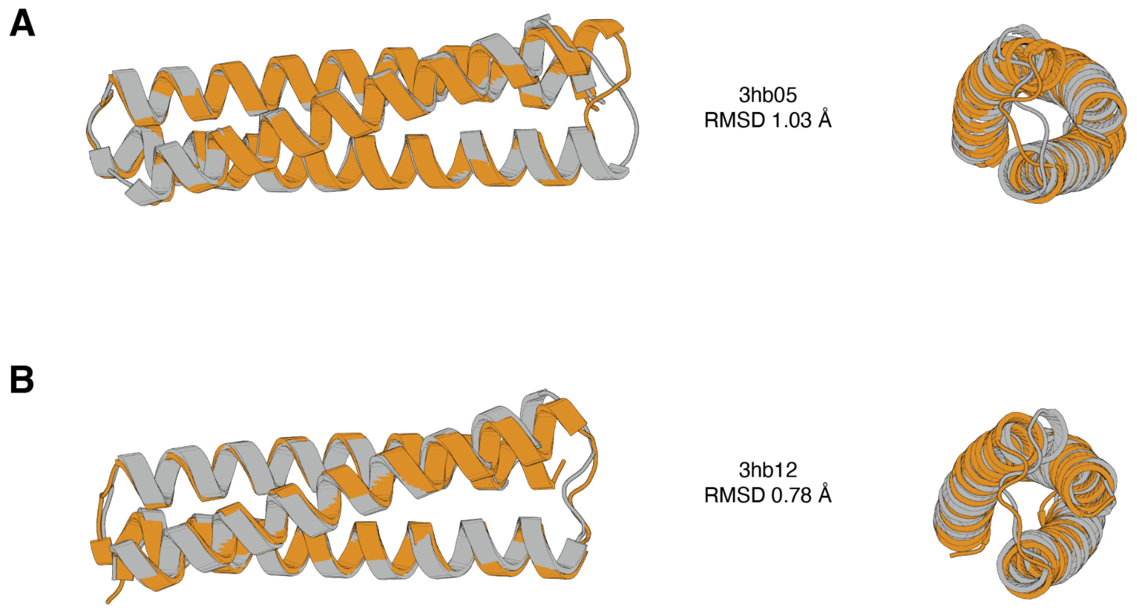


**Figure S13: Structural validation of three-helix bundles. A,B)** Overlay of design model (orange) and crystal structure (gray) in side view (left) and top view (right) for designs 3hb05 (A, also shown in Figure 4E) and 3hb12 (B).

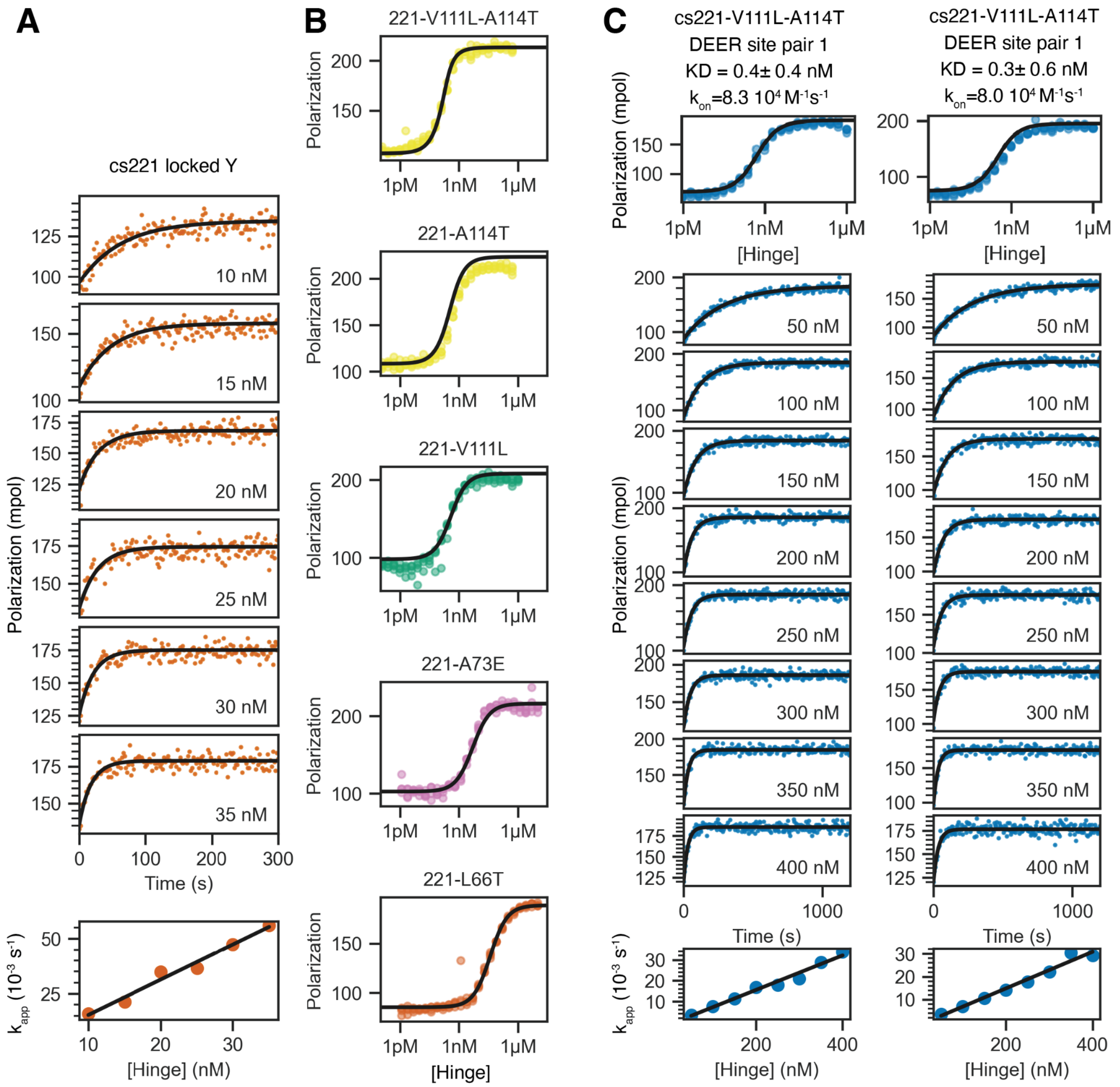


**Figure S14: Additional data supplementing Figure 5. A)** Full FP kinetics experiment for the cs221 locked Y hinge variant shown in Figure 5B. Individual kinetic traces were obtained using 1 nM TAMRA-labeled peptide cs221B and varying hinge concentrations as indicated by plot labels. Single exponential fits give apparent rate constants that increase linearly with the total hinge concentration (orange points in pseudo-first order plot). **B)** Full-range individual plots of the FP titrations shown in Figure 5C. **C)** FP titrations and kinetics for the MTSL-labeled variants of hinge cs221-V111L-A114T. The spin labels have no measurable effect on affinity or association kinetics.

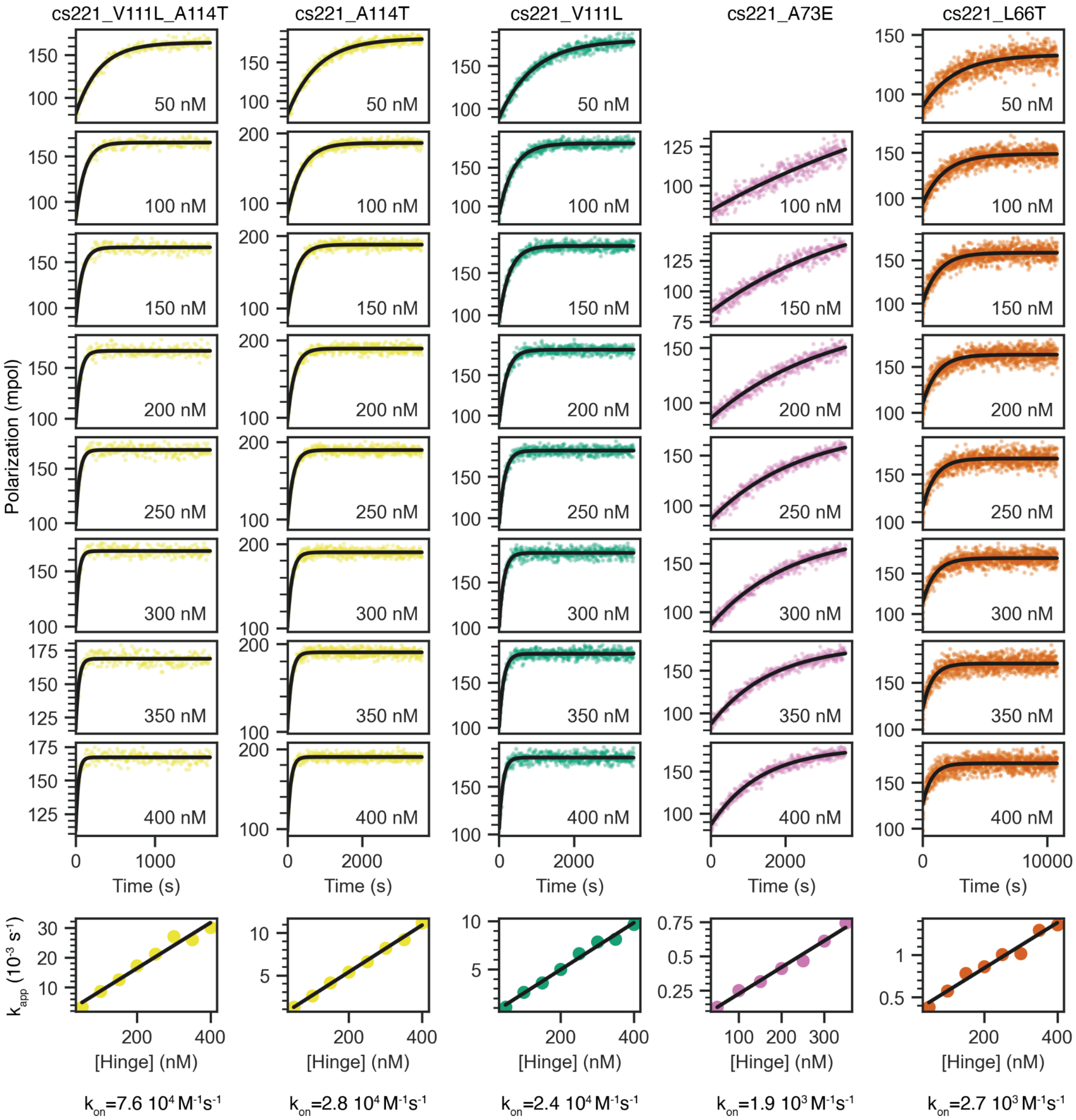


**Figure S15: Full FP kinetics experiments for the cs221 mutants shown in Figure 5C.** Rows 1-8: TAMRA-labeled peptide cs221B at a constant concentration of 5 nM was mixed with hinge at varying concentrations (labels in each plot indicate the hinge concentration for the corresponding experiment). All kinetic traces were fitted using a single-exponential equation (black lines). Row 9: Apparent rate constants from the single exponential fits plotted against the hinge concentration and fitted as linear (black lines). The slope of the linear fit gives the observed on rate $k_{on}$.

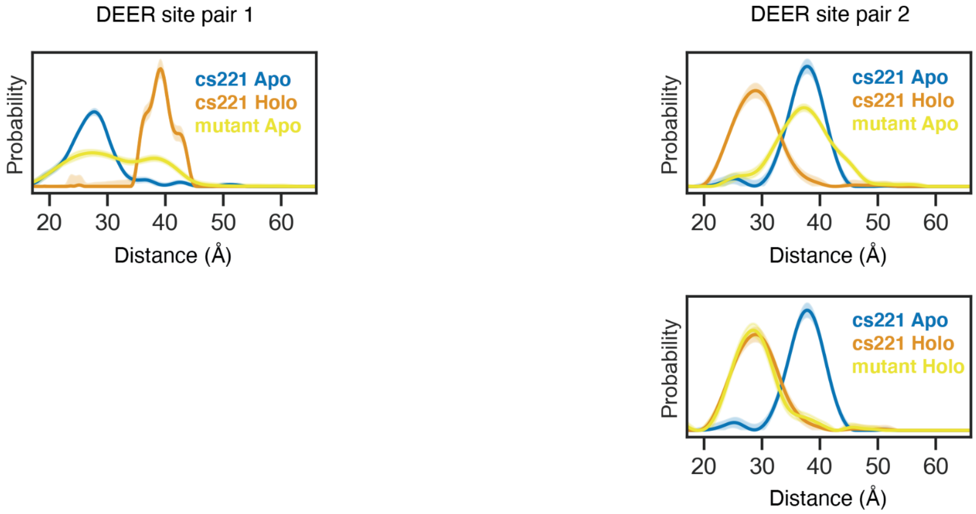


**Figure S16: DEER data on hinge variants that populate both states in absence of peptide.** Distance distributions obtained from DEER experiments with the original hinge cs21 in absence of peptide (blue) and in presence of peptide (orange) as well as of the double mutant cs221-V111L-A114T in absence of peptide (mutant Apo, yellow) and in presence of peptide (mutant Holo, yellow).

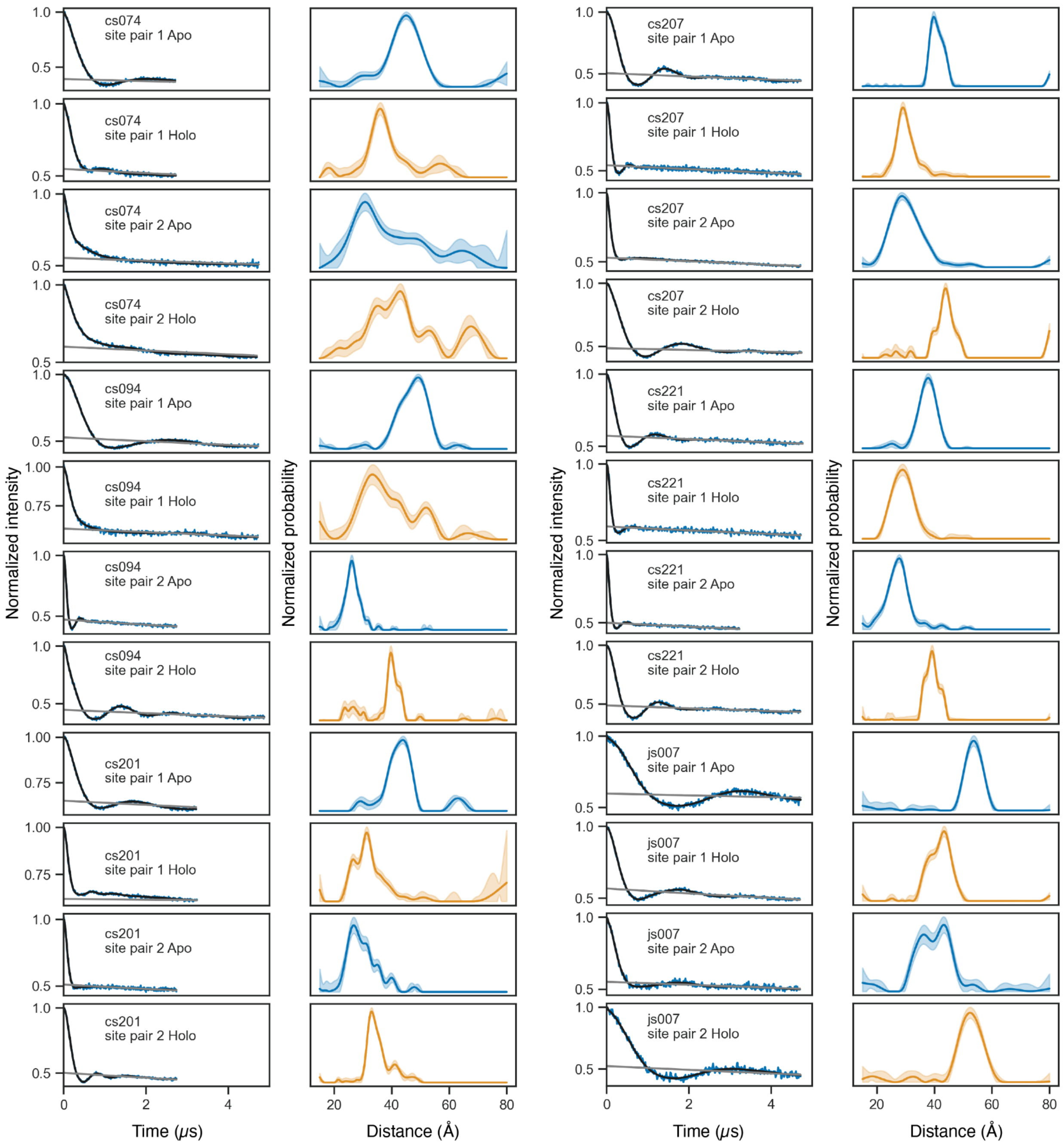


**Figure S17: Additional data for DEER experiments.** Raw DEER traces (blue), foreground fits (black), and background fits (gray) are shown on the left. Distance distributions are shown on the right colored by state (apo: blue, holo: orange) with 95% confidence intervals shown as semi-transparent bands.

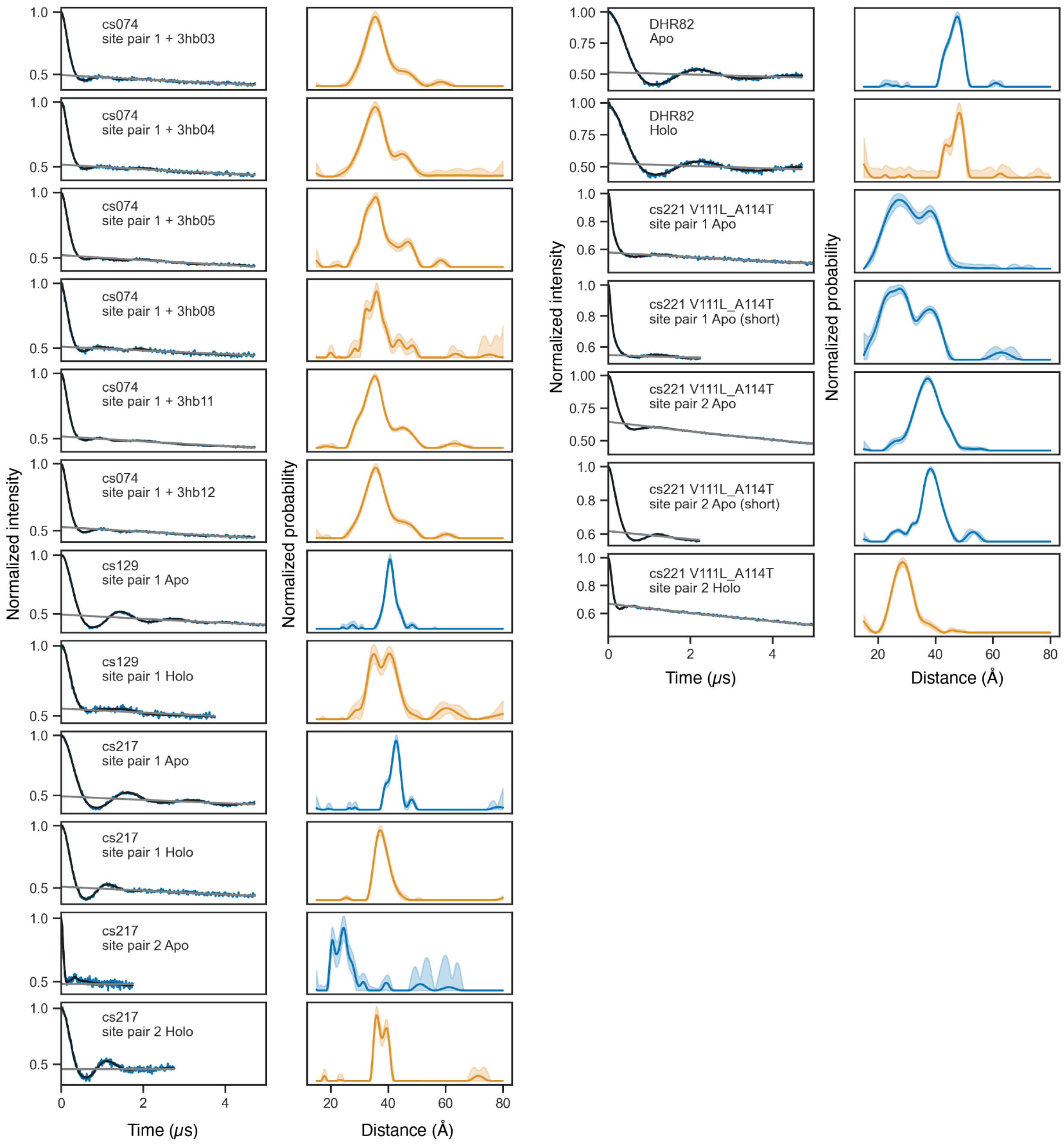


**Figure S18: Additional data for DEER experiments.** Raw DEER traces (blue), foreground fits (black), and background fits (gray) are shown on the left. Distance distributions are shown on the right colored by state (apo: blue, holo: orange) with 95% confidence intervals shown as semi-transparent bands.

**Supplementary Table 1. Crystallographic data collection and refinement**

| | 3hb05 (8FIH) | 3hb12 (8FVT) | cs074AB (8FIT) | cs207A (8FIN) | cs207AB (8FIQ) |
|---|---|---|---|---|---|
| **Data Collection** | | | | | |
| Space group | P 21 21 2 | I 4 | P1 | P 21 21 21 | P 43 21 2 |
| *Cell dimensions* | | | | | |
| *a*, *b*, *c* (Å) | 66.01, 119.75, 38.80 | 67.67, 67.67, 82.37 | 44.11, 45.36, 61.80 | 22.86, 81.73, 154.94 | 71.88, 71.88, 127.64 |
| $\alpha$, $\beta$, $\gamma$ (°) | 90, 90, 90 | 90, 90, 90 | 109.27, 94.54, 104.09 | 90, 90, 90 | 90, 90, 90 |
| Resolution (Å) | 44.35 - 2.2 (2.3 - 2.2) | 47.86 - 3.07 (3.86 - 3.07) | 57.45 - 2.75 (2.9 - 2.75) | 43.66 - 2.3 (2.39 - 2.3) | 47.72 - 2.66 (2.8 - 2.66) |
| $R_{merge}$ | 0.030 (1.161) | 0.098 (0.415) | 0.077 (0.428) | 0.1154 (0.5973) | 0.1361 (4.34) |
| $R_{\mathrm{pim}}$ | 0.030 (0.531) | 0.035 (0.145) | 0.064 (0.369) | 0.03218 (0.1575) | 0.03788 (1.226) |
| $I/\sigma(I)$ | 9.89 (2.16) | 17.39 (5.59) | 7.58 (0.57) | 15.19 (4.77) | 11.84 (0.60) |
| $CC_{1/2}$ | 0.999 (0.852) | 1 (0.978) | 0.986 (0.769) | 1 (0.99) | 1 (0.661) |
| Completeness (%) | 95.27 (72.82) | 99.60 (99.48) | 89.39 (90.08) | 99.85 (99.93) | 97.86 (89.38) |
| Redundancy | 1.4 (1.0) | 8.8 (9.1) | 2.2 (2.3) | 13.7 (14.9) | 13.9 (13.3) |
| **Refinement** | | | | | |
| Resolution (Å) | 44.35 - 2.2 (2.3 - 2.2) | 47.86 - 3.07 (3.86 - 3.07) | 57.45 - 2.75 (2.9 - 2.75) | 43.66 - 2.3 (2.39 - 2.3) | 47.72 - 2.66 (2.8 - 2.66) |
| No. reflections | 15519 (1653) | 30894 (15864) | 10934 (1563) | 13722 (1431) | 9939 (1246) |

| | 3hb05 (8FIH) | 3hb12 (8FVT) | cs074AB (8FIT) | cs207A (8FIN) | cs207AB (8FIQ) |
|---|---|---|---|---|---|
| $R_{work}$ / $R_{free}$ | 0.1908 (0.1892)/ 0.2478 (0.2495) | 0.2248 (0.2998)/ 0.2728 (0.3158) | 0.2457 (0.2901)/ 0.2984 (0.3345) | 0.2403 (0.2360)/ 0.2712 (0.3046) | 0.2691 (0.4680)/ 0.3067 (0.4463) |
| *No. atoms* | | | | | |
| Protein | 2490 | 1518 | 3871 | 2659 | 1538 |
| Water | 81 | 0 | 4 | 24 | 0 |
| Ligand | 10 | 0 | 0 | 0 | 0 |
| Ramachandran Favored/allowed Outlier (%) | 98.70/ 1.30/ 0.00 | 96.81/ 3.19/ 0.00 | 97.49/ 1.88/ 0.00 | 99.42/ 0.58 /0/00 | 99.48/ 0.52/ 0.00 |
| *R.m.s. deviations* | | | | | |
| Bond lengths (Å) | 0.006 | 0.004 | 0.002 | 0.001 | 0.002 |
| Bond angles (°) | 0.560 | 0.590 | 0.440 | 0.300 | 0.380 |
| $B_{factors}$ *(Å$^2$)* | | | | | |
| Protein | 41.25 | 85.94 | 86.88 | 48.07 | 108.65 |
| Water | 38.54 | n/a | 75.76 | 44.70 | n/a |
| Ligand | 64.46 | n/a | n/a | n/a | n/a |

**Supplementary Table 2. DEER experimental metadata and parameters**

| Sample | λ | Scans | Δt (ns) | SNR | $t_0$ offset (ns) | $\tau_2$ (μs) | SRT (ms) | α | β | site pair |
|---|---|---|---|---|---|---|---|---|---|---|
| cs074 D1 Holo | 0.45 | 50 | 11 | 50 | 82.8 | 3 | 1.53 | 0.91 | 0.51 | K36R1 Q211R1 |
| cs074 D1 Apo | 0.57 | 85 | 11 | 67 | 97.2 | 3 | 1.53 | 3.10 | 0.20 | K36R1 Q211R1 |
| cs074 D2 Apo | 0.44 | 31 | 16 | 45 | 73.6 | 5 | 2.04 | 2.50 | 8.60 | E19R1 E179R1 |
| cs074 D2 Holo | 0.40 | 39 | 16 | 79 | 99.2 | 5 | 2.04 | 1.03 | 9.15 | E19R1 E179R1 |
| cs129 D1 Holo | 0.45 | 6 | 12 | 50 | 54 | 4 | 2.04 | 0.62 | 1.13 | E25R1 K176R1 |
| cs129 D1 Apo | 0.50 | 94 | 22 | 85 | 70.4 | 7 | 2.04 | 0.13 | 2.37 | E25R1 K176R1 |
| cs094 D1 Apo | 0.47 | 74 | 16 | 72 | 68.8 | 5 | 2.04 | 1.35 | 1.03 | R30R1 E206R1 |
| cs094 D1 Holo | 0.40 | 83 | 16 | 43 | 80 | 5 | 2.04 | 1.70 | 2.98 | R30R1 E206R1 |
| cs094 D2 Holo | 0.55 | 69 | 16 | 72 | 67.2 | 5 | 2.04 | 0.10 | 1.36 | K12R1 E178R1 |
| cs094 D2 Apo | 0.53 | 49 | 10 | 92 | 68 | 3 | 2.04 | 0.11 | 8.79 | K12R1 E178R1 |
| js007 D1 Apo | 0.45 | 31 | 16 | 40 | 96 | 5 | 2.04 | 1.28 | 6.63 | Q10R1 Q219R1 |
| js007 D1 Holo | 0.48 | 34 | 16 | 31 | 81.6 | 5 | 2.04 | 2.77 | 1.54 | Q10R1 Q219R1 |
| js007 D2 Apo | 0.40 | 40 | 16 | 34 | 78.4 | 5 | 2.04 | 1.51 | 9.24 | D60R1 R190R1 |
| js007 D2 Holo | 0.43 | 43 | 16 | 74 | 97.6 | 5 | 2.04 | 0.45 | 0.38 | D60R1 R190R1 |
| cs207 D2 Holo | 0.46 | 25 | 8 | 56 | 86.4 | 5 | 2.04 | 0.40 | 1.32 | K10R1 D150R1 |
| cs207 D2 Apo | 0.50 | 25 | 16 | 62 | 76.8 | 5 | 2.04 | 0.25 | 0.82 | K10R1 D150R1 |
| cs207 D1 Apo | 0.49 | 85 | 16 | 122 | 84.8 | 5 | 2.04 | 0.24 | 8.68 | D25R1 K130R1 |
| cs207 D1 Holo | 0.50 | 69 | 16 | 99 | 59.2 | 5 | 2.04 | 0.15 | 0.24 | D25R1 K130R1 |
| cs217 D1 Apo | 0.52 | 32 | 6 | 35 | 69.6 | 2 | 2.04 | 0.15 | 0.18 | E47R1 R127R1 |
| cs217 D1 Holo | 0.54 | 17 | 10 | 37 | 47 | 3 | 2.04 | 0.13 | 6.17 | E47R1 R127R1 |
| cs217 D2 Apo | 0.51 | 63 | 16 | 91 | 84.8 | 5 | 2.04 | 0.15 | 8.61 | R27R1 E147R1 |
| cs217 D2 Holo | 0.49 | 74 | 16 | 57 | 72 | 5 | 2.04 | 0.48 | 8.91 | R27R1 E147R1 |
| cs217 D1 Apo | 0.52 | 110 | 8 | 35 | 69.6 | 3 | 2.04 | 0.15 | 0.18 | E47R1 R127R1 |
| cs221 D1 Apo | 0.50 | 69 | 10 | 98 | 82 | 3.5 | 2.04 | 0.02 | 0.63 | E43R1 K131R1 |
| cs221 D1 Holo | 0.52 | 72 | 16 | 74 | 88 | 5 | 2.04 | 0.12 | 0.24 | E43R1 K131R1 |
| cs221 D2 Apo | 0.43 | 47 | 16 | 62 | 76.8 | 5 | 2.04 | 0.86 | 0.25 | R23R1 E150R1 |
| cs221 D2 Holo | 0.41 | 48 | 16 | 39 | 76.8 | 5 | 2.04 | 1.50 | 9.21 | R23R1 E150R1 |

| Sample | λ | Scans | Δt (ns) | SNR | $t_0$ offset (ns) | $\tau_2$ (μs) | SRT (ms) | α | β | site pair |
|---|---|---|---|---|---|---|---|---|---|---|
| cs201 D1 Holo | 0.50 | 83 | 8 | 154 | 76 | 3 | 2.04 | 0.14 | 5.42 | R35R1 E207R1 |
| cs201 D1 Apo | 0.49 | 9 | 8 | 61 | 77.6 | 3 | 2.04 | 0.46 | 9.16 | R35R1 E207R1 |
| cs201 D2 Apo | 0.35 | 55 | 12 | 91 | 90 | 3.5 | 2.04 | 0.42 | 9.23 | K24R1 E180R1 |
| cs201 D2 Holo | 0.36 | 90 | 12 | 86 | 76.8 | 3.5 | 2.04 | 0.18 | 2.93 | K24R1 E180R1 |
| DHR82P D1 Apo | 0.50 | 238 | 16 | 66 | 81.6 | 5 | 2.04 | 0.60 | 8.80 | K36R1 A211R1 |
| DHR82P D1 Holo | 0.49 | 92 | 16 | 43 | 86.4 | 5 | 2.04 | 0.37 | 0.46 | K36R1 A211R1 |
| cs074 3hb03 D1 Holo | 0.51 | 78 | 16 | 75 | 73.6 | 5 | 2.04 | 1.17 | 0.60 | K36R1 Q211R1 |
| cs074 3hb04 D1 Holo | 0.49 | 21 | 16 | 75 | 81.6 | 5 | 2.04 | 1.18 | 1.03 | K36R1 Q211R1 |
| cs074 3hb05 D1 Holo | 0.50 | 60 | 16 | 106 | 83.2 | 5 | 2.04 | 0.37 | 8.76 | K36R1 Q211R1 |
| cs074 3hb08 D1 Holo | 0.49 | 21 | 16 | 63 | 83.2 | 5 | 2.04 | 0.19 | 1.19 | K36R1 Q211R1 |
| cs074 3hb11 D1 Holo | 0.48 | 236 | 16 | 192 | 80 | 5 | 2.04 | 0.34 | 0.00 | K36R1 Q211R1 |
| cs074 3hb12 D1 Holo | 0.47 | 73 | 16 | 102 | 84.8 | 5 | 2.04 | 0.77 | 3.49 | K36R1 Q211R1 |
| cs221-mut D1 Apo | 0.42 | 158 | 18 | 66 | 81 | 7 | 2.04 | 1.69 | 8.68 | E43R1 K131R1 |
| cs221-mut D2 Apo | 0.36 | 186 | 18 | 151 | 84.6 | 7 | 2.04 | 0.57 | 9.16 | R23R1 E150R1 |
| cs221-mut D2 Holo | 0.41 | 48 | 16 | 43 | 76.8 | 5 | 2.04 | 1.55 | 6.76 | R23R1 E150R1 |

**λ - Modulation depth**
**Δt - Pump pulse time step**
**SRT - Shot repetition time**
**α - Smoothness regularization parameter**
**β - Compactness regularization parameter**
